# Development and translational validation of a novel mouse model for allergic rhinitis

**DOI:** 10.64898/2026.09.03.749032

**Authors:** Lubnaa Hossenbaccus, Wil P. Taylor, Sara Teimouri Nezhad, Jack M. Taylor, Cortney Haird, Andisheh Liaghat, Sarah E. Hopkins, Vidthiya Jeyanathan, Tyson Rudolf, Adrian J. Paz, Hannah Botting, Lisa M. Steacy, Fazila Chouiali, Terry Walker, Joaquin Sanz, Sarah Garvey, Jan Schinköthe, Anne K. Ellis, Eva Kaufmann

## Abstract

**Background:** Approximately 30% of the North American population are affected by seasonal or perennial rhinitis. Allergens from cats are common triggers of allergic rhinitis. Importantly, cat allergic rhinitis (cat-AR) is a major risk factor for more severe allergic diseases, including asthma. Current therapies rarely achieve full symptom control and development of novel therapeutics is constrained by the lack of physiologically relevant preclinical models.

**Objective:** Establishment of a clinically-informed mouse model of cat allergen exposure in humans.

**Methods:** Controlled human exposure was performed using airborne natural cat dander (CD) in a validated environmental exposure facility. Nasal immune cells and IgE concentrations in serum and nasal samples were assessed. In mice, we established a 14-day intermittent intranasal sensitization and challenge protocol using the same CD allergens. Clinical signs in mice were objectively quantified using AI-based video detection. Cellular changes in the nose were assessed with level-specific, spatial resolution. Lastly, systemic sensitization was confirmed through serum IgE and intradermal ear challenge.

**Measurements and Main Results:** Following CD exposure, cat-allergic participants displayed increased nasal symptoms and eosinophil influx. CD-sensitized mice recapitulated key clinical features, including increased nasal rubbing, elevated nasal immune cell infiltration, and systemic sensitization. Importantly, spatial investigation of the cellular changes revealed major changes in the turbinate and olfactory regions of the nose.

**Conclusions:** This novel mouse model provides a translationally valuable platform for investigating local and systemic mechanisms of cat-AR. It will serve for mechanistic and therapeutic evaluations with the goal of improving clinical translation and therapeutic precision.

## INTRODUCTION

Allergic rhinitis (AR) affects between 20-40% of the North American population (1, 2). Cat allergens are among the most common triggers of perennial allergies, with >20% of individuals sensitized (3). 8 *Felis domesticus* (Feld) cat allergens have been identified. Most cat-AR patients are sensitized to Feld1 (4). Feld1 is a small protein of approximately ¼ of the size of pollen grains (5µm vs 20µm) (5). The small size and sticky nature of Feld1 facilitates its dissemination through clothing and in indoor environments (6) which results in widespread and persistent exposure. Clinically, cat-AR presents with impaired breathing, sneezing, nasal congestion or rhinorrhea, and often itchy, watery eyes (7). Beyond these symptoms, cat-AR substantially impacts quality of life through reduced productivity and sleep and imposes a significant economic burden, estimated at $3.4 billion annually in the USA (8). Importantly, cat-AR is a risk factor for more severe, potentially life-threatening allergic diseases: cat-AR is specifically associated with 2x increased probability for asthma development (9, 10). Asthma is a major driver of healthcare burden (>$25 billion annually) (11). As the prevalence of allergic diseases continues to rise globally, the burden of cat-AR and its associated risks is likewise expected to increase.

Management of cat-AR is complicated by the ubiquitous and persistent nature of cat allergens. Due to their adhesive properties, allergen levels remain elevated for weeks even after removal of the source (12), and detectable Feld1 is present in the majority of indoor environments, including those without cats (13, 14). Thus, avoidance of cat allergens, as most other allergic rhinitis triggers, is often impossible.

Current pharmacologic management of allergic rhinitis follows a stepwise approach based on disease severity, including antihistamines, intranasal corticosteroids, leukotriene receptor antagonists, and allergen immunotherapy (AIT). For cat-AR, subcutaneous immunotherapy (SCIT) remains the only approved AIT modality (15). SCIT is safe and can reduce symptom severity (16) but is mainly standardized to Feld1 (15) and rarely achieves full symptom resolution (17). Although >90% of patients are sensitized to Feld1, only ∼27% are mono-sensitized (18).

Sensitization to Feld2/4/5 occurs in 40–60% of patients (18). Improved therapeutic strategies that better address the complex clinical picture are urgently needed. Novel therapeutics for cat-AR are currently in active development (19–21). Advancing such candidates would profit from experimental models that recapitulate the physiological context of human allergen exposure. While *in vitro* systems, including human nasal epithelial cultures, provide valuable mechanistic insight, they do not capture systemic immune responses or the dynamic recruitment of immune cells to the nasal mucosa. Small animal models are therefore essential for studying disease pathogenesis and evaluating therapeutic interventions in an integrated system.

Several murine models of cat-AR have been described (**Table S1**). All of them rely on subcutaneous or intraperitoneal allergen administration, often combined with adjuvants as nonspecific immune enhancers. Although informative for studying allergen-responsive pathways, these approaches do not reflect the natural route, composition, or kinetics of cat allergen exposure in humans. In contrast, real-life sensitization occurs primarily through repeated inhalation of airborne allergens in complex environmental compositions (22). Preclinical models that recapitulate physiological exposure remain lacking.

Guided by a controlled human allergen exposure model, we here developed a clinically-informed murine model of cat-AR that incorporates natural cat dander (CD) exposure via the nasal mucosa. By aligning exposure conditions and outcome measures between species, this model demonstrates key features of human cat-AR and provides a platform for investigating disease mechanisms and evaluating novel therapeutic strategies with improved translational relevance.

## METHODS

Methodological details are available in the online supplement.

### Cat dander allergen

Cat dander was obtained from Stallergenes Greer, USA (RME63P).

### Human clinical study

Protocols and procedures were approved by Queen’s University Health Sciences and Affiliated Teaching Hospitals Research Ethics Board as published (23). 31 confirmed cat-allergic participants with ≥2-year clinical history of cat-allergy symptoms and positive skin prick test to cat hair, and 15 non-allergic participants with no history of cat allergies and a negative skin prick test to all tested allergens (12-65 years old) were included in this study and provided informed consent. Participants were exposed to airborne CD in the SPaC-EEU for 3h. Air samples revealed mean Feld1 exposure concentration of 69ng/m^3^. Symptom scores and biological samples were collected at various timepoints (**Table S3**).

### Mouse experiments

#### Mice

Male and female 7-12-week-old BALB/c and C57BL/6 mice were purchased from Jackson Laboratories or bred at Queen’s University Animal Facility, or the Animal Care Service Facility of the Research Institute of the McGill University Health Centre (RI-MUHC). Mice were housed under specific pathogen-free conditions. All animal studies were approved by the Animal Research Ethics Council at Queen’s University (protocol 2338) and McGill University (protocol MUHC- 10263).

#### Mouse model of cat allergic rhinitis

Aligned with the human exposure level, mice were intranasally (IN) exposed to 54ng of Feld1 (days 1/2/3; 7/8/9) in 9µL of reconstituted CD under isoflurane anesthesia. No-treatment-control mice underwent anesthesia with no IN exposure, whereas saline-control mice received anesthesia and IN sterile saline. Mice were monitored for 30min after each administration. All mouse experiments were performed in the morning. Mice were euthanized by CO2 asphyxiation under isoflurane anesthesia.

#### Distribution of fluid following intranasal administration

To delineate the anatomical extent of fluid inhalation in IN allergen exposure, 9µL of Evan’s Blue solution (0.125% w/v in saline) was administered to anesthetized mice. After 5min and once fully awakened, mice were anesthetized again and directly euthanized by CO2 asphyxiation.

#### Intradermal ear challenge

For intradermal ear challenge, sterile saline was injected intradermally in the left ear, and CD containing 54ng Feld1 in the right ear, using a 31G needle. Ear thickness measurements were collected using a digital micrometer (Beslands, accuracy 0.003 mm).

#### Mouse serum isolation and IgE quantification

Blood samples were collected from anesthetized mice via cardiac puncture into lithium heparin tubes (BD). Total IgE in mouse serum samples was determined using ELISA MAX™ Standard Set Mouse IgE (Biolegend).

#### Nasal fluid collection method

A 22G feeding tube on a 10mL syringe filled with 10mL sterile saline was inserted into the nasopharyngeal sphincter. The plunger was slowly depressed to flush the nasal cavity, avoiding spillover that would rinse the oral side of the soft and hard palate. The flushing of the nasal cavity was repeated once using a 20G feeding tube.

#### Flow cytometry

Flow cytometry on nasal cells was performed according to standard procedure, details are provided in the supplemental methods.

#### Histology

Skulls were fixed in 4% formaldehyde for 24h. Decalcification, histological and immunohistochemical staining were performed according to standard protocols. Slides were analyzed by a blinded, board-certified veterinary pathologist (Dipl. ACVP), using QuPath (version 6.0.0)(24).

#### AI-assisted behavioural analysis of nasal rubbing in mice

Mice were video recorded in their home cages for 30min following each exposure. To enable objective and blinded behavioural quantification, we developed an AI-powered video analysis and annotation tool to identify candidate nasal rubbing events. All candidate events were manually reviewed and classified by a blinded researcher to distinguish nasal rubbing from grooming, feeding, or unrelated movements. An episode of nasal rubbing was characterized by the starting of nasal rubbing behaviour until termination, independent of the specific number of individual rubs on the nose.

### Statistical Analysis

Statistical analyses were performed using Graph Pad Prism Version 10. Data are displayed as mean±SEM. Statistical significance was determined as indicated in the figure legends.

## RESULTS

### Experimental allergen exposure elicits allergic responses in cat-sensitized patients

Thirty-one cat-allergic participants and 15 healthy, non-allergic controls provided informed consent and were exposed for 3h to an average of 69ng/m^3^ of airborne cat allergens in the SPaC- EEU (**Fig 1A**), reflective of exposure in both homes with and without cats (25, 26) (**Table S2**).

**Fig 1.**
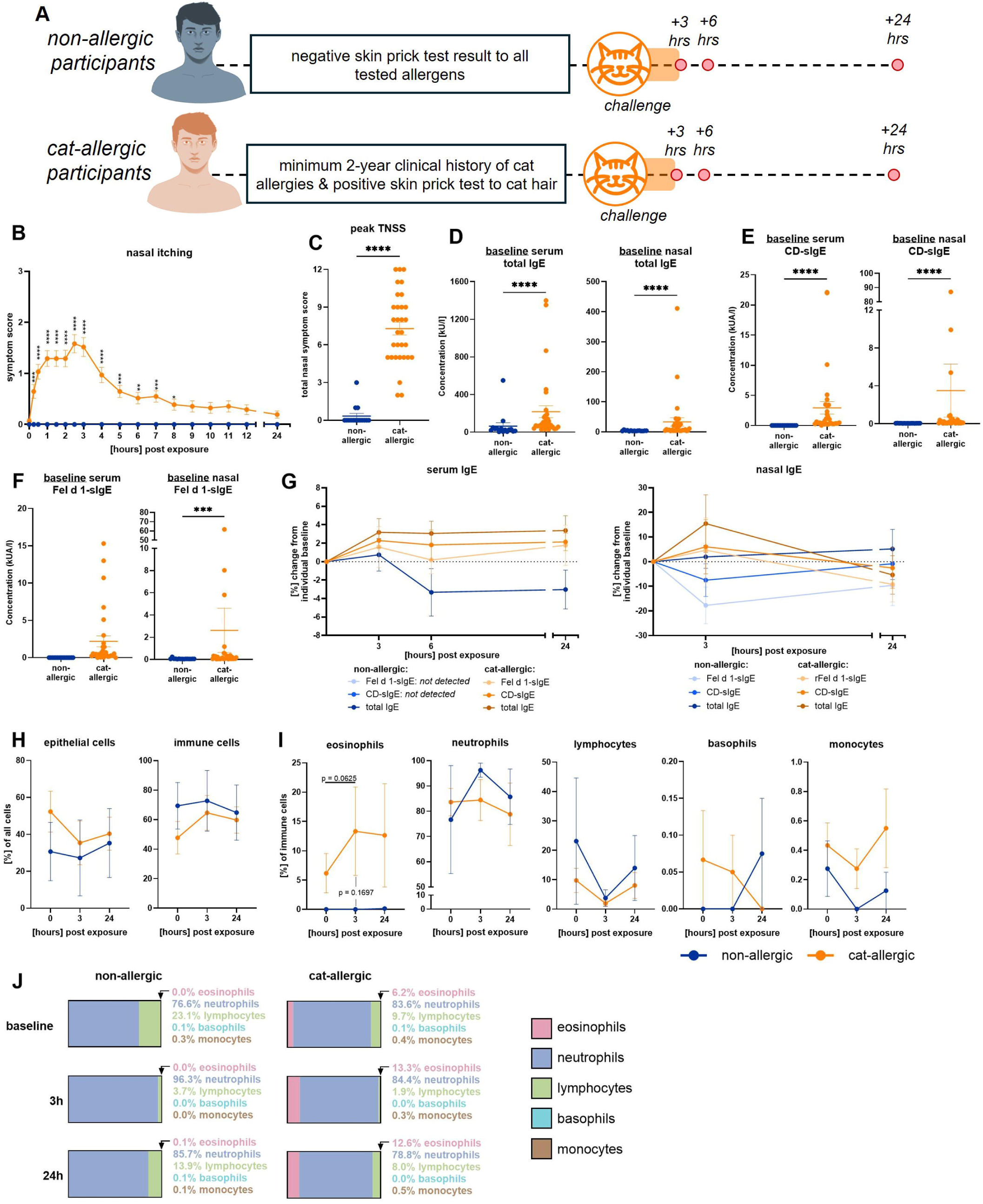
Clinical exposure in the SPaC-EEU elicited increased nasal symptoms and nasal eosinophils in cat-allergic participants. **(A)** Experimental model for exposure of cat-allergic (n=31) and non-allergic (n=15) participants to airborne CD in the SPaC-EEU. **(B)** Kinetic of nasal itching scores from onset of CD exposure. **(C)** Peak Total Nasal Symptom Scores (TNSS) at 2.5h post CD exposure onset. **(D)** Baseline total IgE levels in serum (left panel) and nose (right panel). **(E)** Baseline CD-specific IgE (CD-sIgE) levels in serum (left panel) and nose (right panel) **(F)** Baseline Feld1-sIgE levels in serum (left panel) and nose (right panel). **(G)** Development of serum (left panel) and nasal (right panel) IgE levels post CD exposure. **(H-J)** Nasal lavage sampling of n=10 cat-allergic and n=4 non-allergic participants. Assessment of **(H)** epithelial and **(I, J)** immune cell numbers in nasal lavage samples. Mann-Whitney test. Mixed-effects model with Sidak’s multiple comparisons test. *p≤0.05, **p<0.01, ***p<0.001, ****p<0.0001.

Cat-allergic participants experienced significantly elevated symptoms of nasal itching and sneezing, starting from 15min after exposure onset and lasting for up to 8h (**Fig 1B**, **S1A**), as well as rhinorrhea/post-nasal drip and nasal congestion/stuffiness for the entire study period (**Figure S1B-C**) Summing these individual symptoms as a Total Nasal Symptom Score (TNSS), cat-allergic participants had a peak in TNSS at 2.5h post exposure onset, which was significantly elevated compared to non-allergic participants (**Fig 1C**).

At baseline, cat-allergic participants had significantly higher concentrations of total IgE (**Fig 1D**), cat dander-specific IgE (CD-sIgE) (**Fig 1E**), and Feld1-sIgE (**Fig 1F**) in both serum and nasal samples. After 3h of CD exposure, serum IgE levels increased by ∼3% only in cat-allergic participants, and nasal IgE levels increased by ∼15% (**Fig 1G**).

CD exposure led to non-significant changes in epithelial and immune cell counts in the nasal lavages of non-allergic and cat-allergic participants (**Fig 1H, S2**). However, shifts in immune cell composition occurred, with substantial inter-individual variability. (**Fig 1I-J**). In all participants, both at baseline and after CD exposure, neutrophils predominated (76.6-83.6%) (**Fig 1I-J**). Only in cat-allergic participants, eosinophils were present at baseline (6.2% of immune cells) (**Fig 1I-J**). The most prominent increase in nasal immune cells for cat-allergic participants occurred in eosinophils after 3h of CD exposure (to 13.3% of immune cells, p=0.06, (**Fig 1I**), and eosinophils remained elevated at 24h post-exposure (12.6% of immune cells). Eosinophil accumulation post exposure did not occur in non-allergic participants (**Fig 1I-J**).

### Development of a clinically informed mouse model of cat allergic rhinitis

We used the experimental exposure data from human participants to develop a pre-clinical model of cat-AR. Based on an average inhalation rate of 6L/min (27), a 3h exposure in the SPaC-EEU to ∼50ng/m³ CD corresponds to an estimated human Feld1 exposure of ∼54ng. Hence, we administered 54ng of Feld1 in 9µl saline to isoflurane-anesthetized mice (**Fig 2A**). To determine the distribution upon administration, we traced the same volume of Evan’s Blue solution. We did not observe Evan’s Blue staining past the larynx, which confirmed that an IN administered volume of 9µL localized the exposure to the upper airways, i.e., nares, nasal cavity, and sinuses (**Fig 2B**).

**Fig 2.**
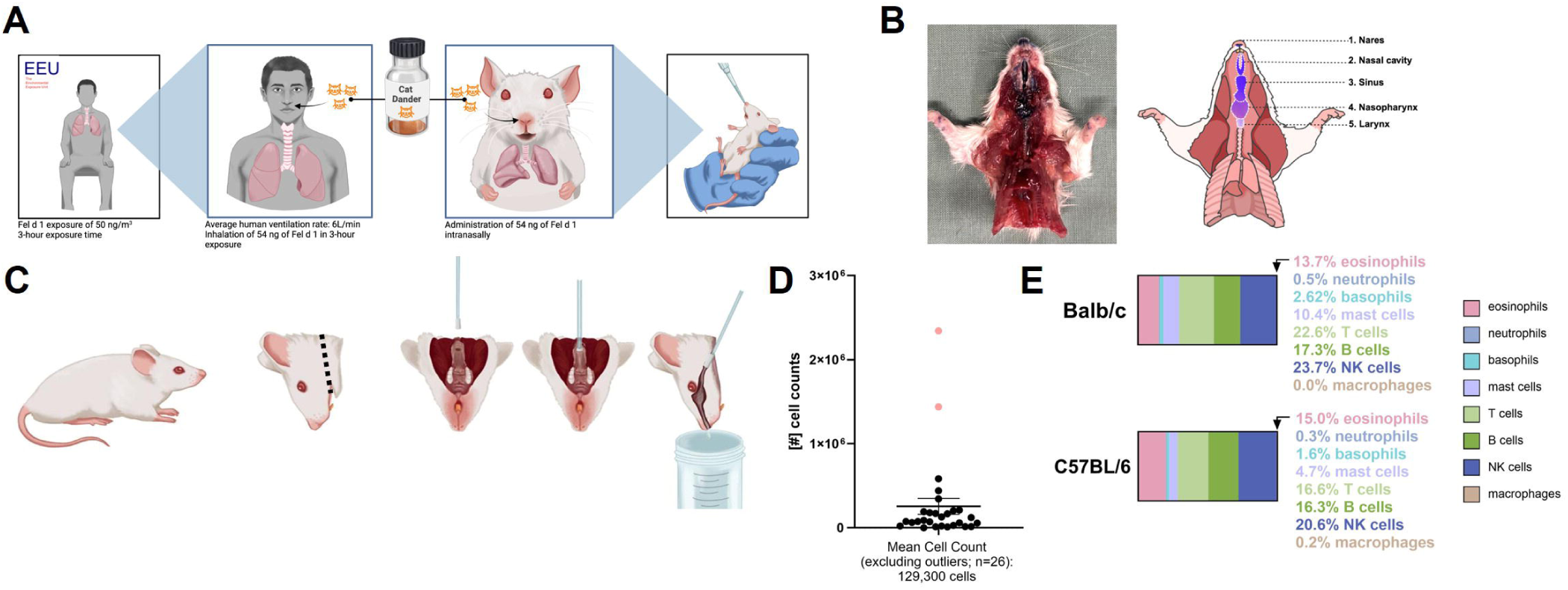
Development of a clinically informed mouse model of cat allergic rhinitis. **(A)** Mice were intranasally administered 54 ng of Feld1, equivalent to the average amount a human adult inhaled over a 3-hour exposure period in the SPaC-EEU. **(B)** Mice were administered 9µL Evan’s Blue solution to determine the extent of the IN distribution in the respiratory track. Representative image and scheme. **(C)** Sampling methodology for nasal immune cells in mice. **(D)** Cell counts from nasal samples with no blood contamination (black dots) and excluded outliers (red dots) (n=26 naïve mice). **(E)** Baseline nasal immune cell composition in BALB/c and C57BL/6 mice (n=3/group).

We optimized a nasal sampling methodology in mice, mimicking the human nasal lavage collection method (**Fig 2C**). Flushing the nasal cavity through the nasopharyngeal sphincter, enabled capture of a substantial number of immune cells from the nasal mucosa (mean nasal cell count of 129,300 cells, **Fig 2D**) without causing bleeding and contaminating the samples with blood cells. Sample points contaminated by blood cells (red marks, **Fig 2D**) could be determined as outliers (defined as +/-2 standard deviations) and were excluded from analysis.

To assess the impact of genetic background on nasal immune cell composition, we compared immune cells in the nasal cavity of C57BL/6 and BALB/c mice. We confirmed equivalent baseline profiles (**Fig 2E**). Considering that genetics are an important risk factor for allergic rhinitis in humans, and we observed no baseline differences in nasal immune cells between mouse strains, we focused subsequent experiments on type-2-biased BALB/c mice.

### Intermittent intranasal cat dander exposure elicited clinical signs of allergic rhinitis in mice

Following the assessment of a suitable administration volume, we implemented a 14-day intermittent CD sensitization and challenge schedule (**Fig 3A**). Control mice received either saline, to account for saline effects, or no administration, only isoflurane, to account for volume effects. Appearance, behaviour, and clinical signs were recorded for 30min after every administration timepoint. No group showed any weight loss (**Fig 3B**), declines in outer appearance, based on posture, coat, and eyes (**Fig 3C**), or activity, i.e., alertness and responsiveness (**Fig 3D**), over the duration of the experiment.

**Fig 3.**
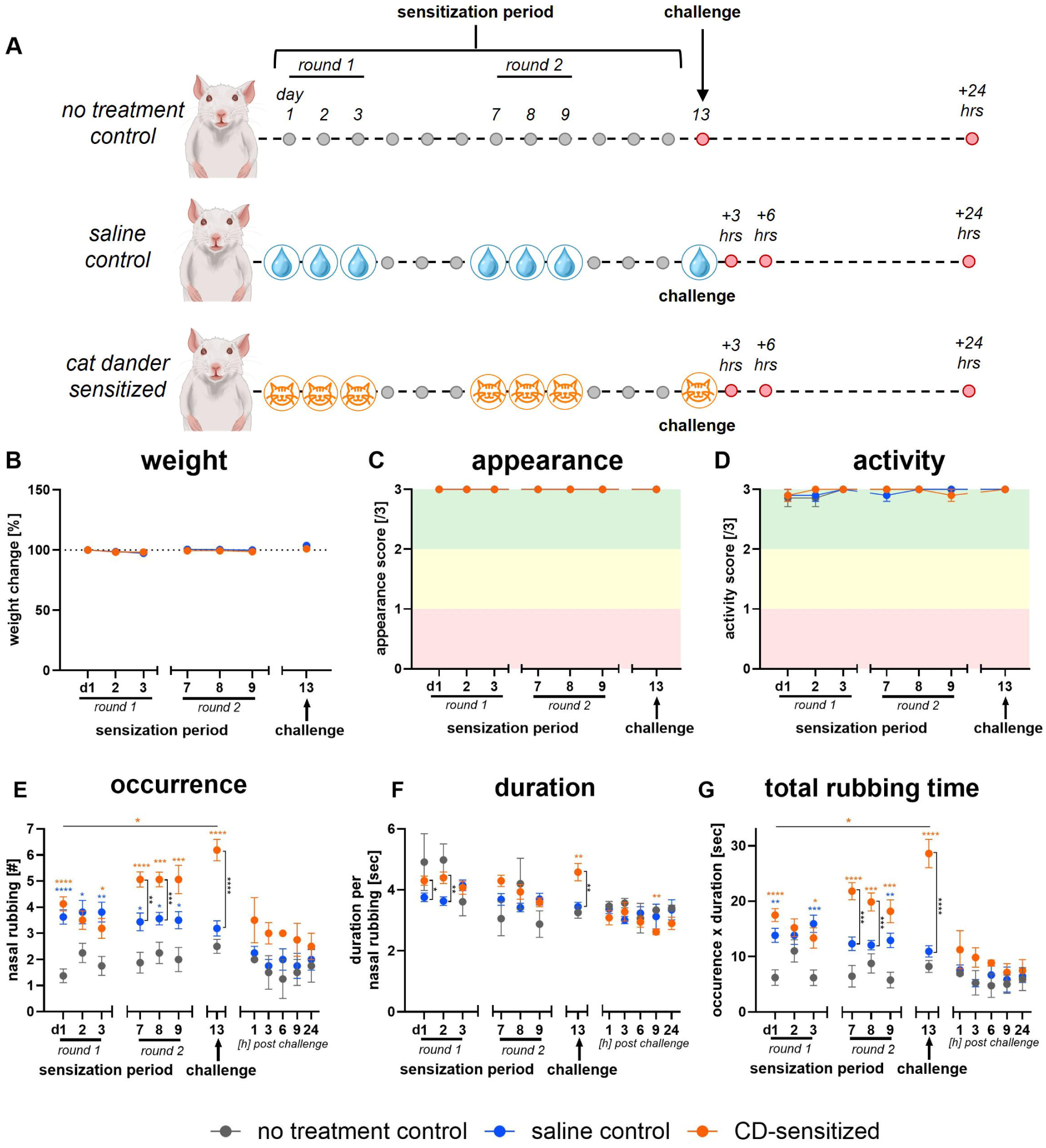
Cat dander exposure induced localized clinical signs in mice. **(A)** Experimental model for 14-day intermittent sensitization and challenge schedule. **(B)** Weight development. **(C)** Appearance. **(D)** Activity in mice (n=16-28/group). Background colours indicate clinical severity assessment. **(E)** Occurrences of nasal rubbing episodes during the first 30min post exposure (n=8-16/group). **(F)** Mean duration of nasal rubbing episodes. **(G)** Total rubbing time calculated as occurrence number x duration. Mixed-effects analysis with Tukey’s multiple comparisons test. *p≤0.05, **p<0.01, ***p<0.001, ****p<0.0001. Orange stars: comparison between CD-sensitized and no treatment control; blue stars: comparison between saline control and no treatment control.

We developed a code for segregation of occurrences of nasal rubbing episodes (**Figure S3**). Both saline-control and CD-sensitized mice increased nasal rubbing episodes compared to no-treatment-controls after day 1 exposure (**Fig 3E**). Following the allergen challenge on day 13, CD-sensitized mice had significantly more instances of nasal rubbing in comparison to days 1-3 and relative to both saline- and no-treatment-control mice (**Fig 3E**). Duration of nasal rubbing was significantly increased for CD-sensitized mice compared to saline- and no-treatment-controls following the challenge (**Fig 3F**). Taking both occurrence and duration into consideration, total rubbing time was significantly higher for CD-sensitized mice on the challenge day compared to sensitization day 1, and compared to both saline- and no-treatment-controls (**Fig 3G**).

### Nasal cat dander exposure stimulates systemic sensitization

We evaluated whether CD exposure initiated systemic sensitization, measured as total IgE concentration in serum. CD-sensitized mice had significantly elevated levels of total IgE at 3h post-IN challenge (**Fig 4A**).

**Fig 4.**
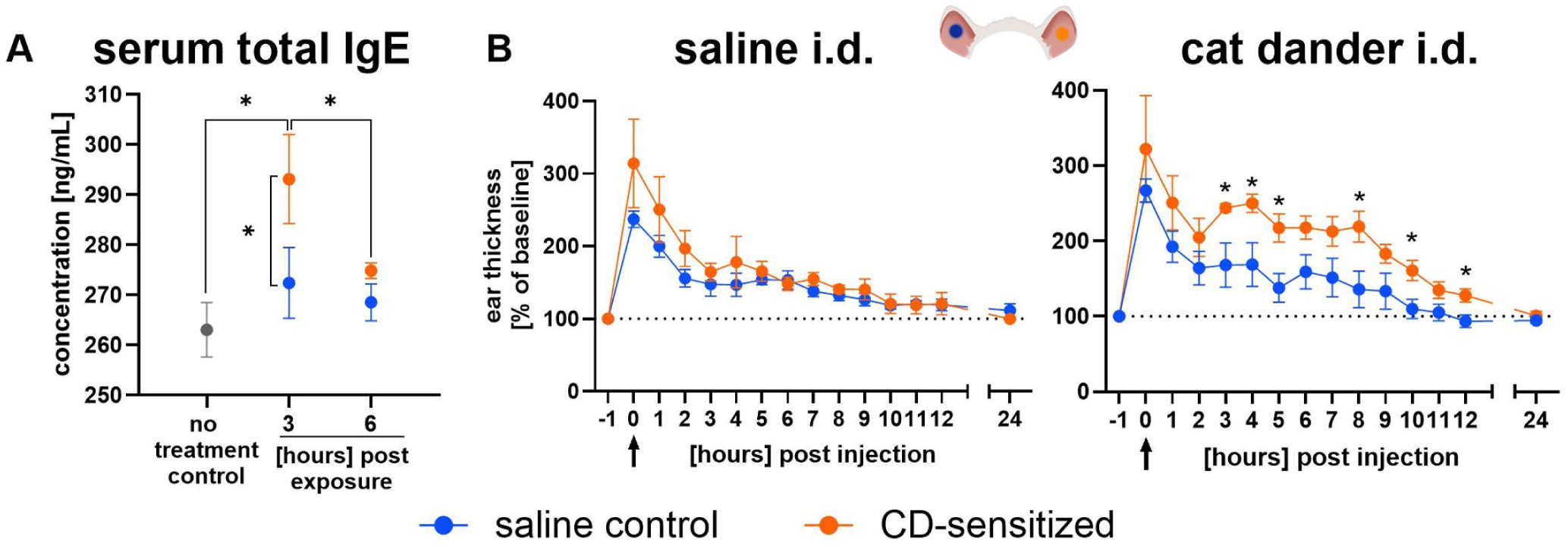
Intranasal cat dander exposure stimulates systemic sensitization. **(A)** Total IgE levels in serum (n=5-8/group). **(B)** Ear thickness in response to intradermal (i.d.) saline (left) and cat dander extract injection (right) (n=4 mice/group). Unpaired t tests. Mann-Whitney tests. *p≤0.05.

Systemic sensitization led to functional effects at locations distant from the nasal cavity, as detected in intradermal ear challenge. Bolus injection increased ear thickness by 200-300% due to volume effect, that gradually decreased over 24h (**Fig 4B**). However, previously CD-sensitized mice experienced a significant secondary response only to CD injection, beginning at 3h post-injection, whereby ear thickness remained significantly increased for 12h (**Fig 4B**).

### Cat dander-sensitized mice have increased eosinophils in the nose at 6 hours post exposure

After confirming systemic sensitization, we determined immune cell changes occurring locally in the nose. Eosinophils were increased in the nasal lavage of both saline-control (19.2%) and CD-sensitized mice (19.7%) at 3h post-challenge compared to no-treatment-control mice (16.9%) **(Fig 5B-C**). Only in CD-sensitized mice, nasal eosinophils further increased at 6h post challenge (21.6%) (**Fig 5B-C**) and were significantly elevated in total cell numbers (**Figure S4**).

**Fig 5.**
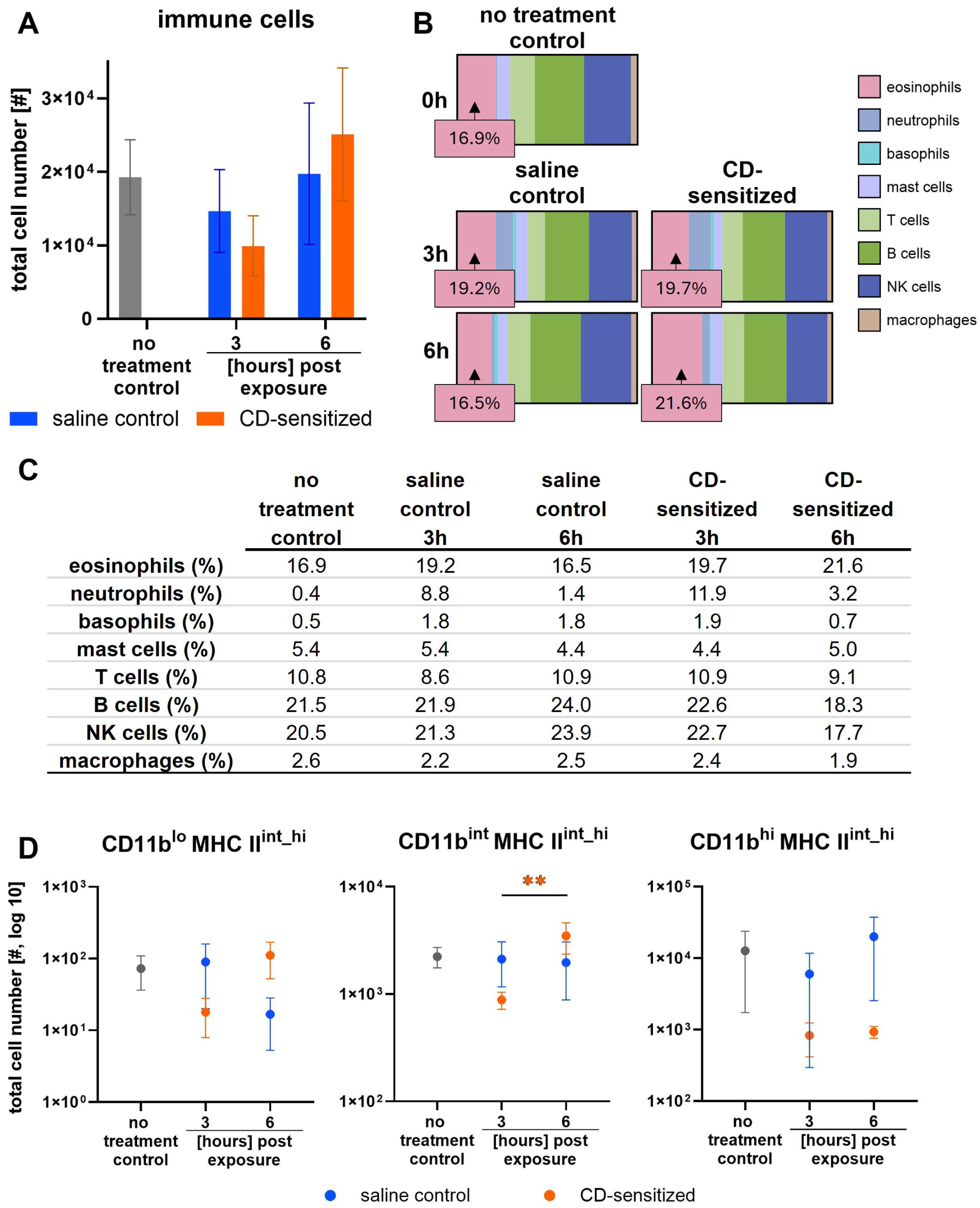
Cat dander-sensitized mice have increased eosinophils in the nose at 6 hours post-challenge. **(A)** Total CD45^+^ immune cell numbers in the nose (n=8 mice/group) **(B, C)** Frequencies of immune cells in the nose (n=8 mice/group). **(D)** Total cell numbers of CD11b^lo^, CD11b^int^, and CD11b^hi^ MHC II^int_hi^ eosinophils in the nose. Mann-Whitney tests. **p<0.01.

Most eosinophils in the nasal mucosa had low or intermediate CD11b expression, in accordance with new recruitment (28), and intermediate or high MHC II expression indicating cellular activation and antigen presentation (29) (**Fig 5D, S5**). Particularly, CD11b^int^MHC II^int_hi^ eosinophil numbers increased in the nasal mucosa of CD-sensitized mice at 6h post-challenge (**Fig 5D**).

### Immune cell numbers increased specifically in the turbinate region of the nose

Beyond the increase in immune cells that was detected by flowcytometry in the nasal lavage fluids, we determined the localization of these immune cells in the nasal mucosa. Therefore, skulls were sampled 3h post challenge and were sectioned at distinct nasal levels: Level 1 (nasal passages), Level II (naso- and maxillo-turbinates), and Level III (olfactory portion of the nasal cavity) (30) (**Fig 6A**). Quantification of CD45-immunopositive cells revealed equal numbers in nasal passage and turbinates in saline-control mice, with lower numbers in olfactory region (**Fig 6B, C**). CD45-immunopositive cells were significantly increased in turbinate and olfactory regions compared to the nasal passage in CD-sensitized mice, with highest numbers in the turbinates (**Fig 6B-C**).

**Fig 6.**
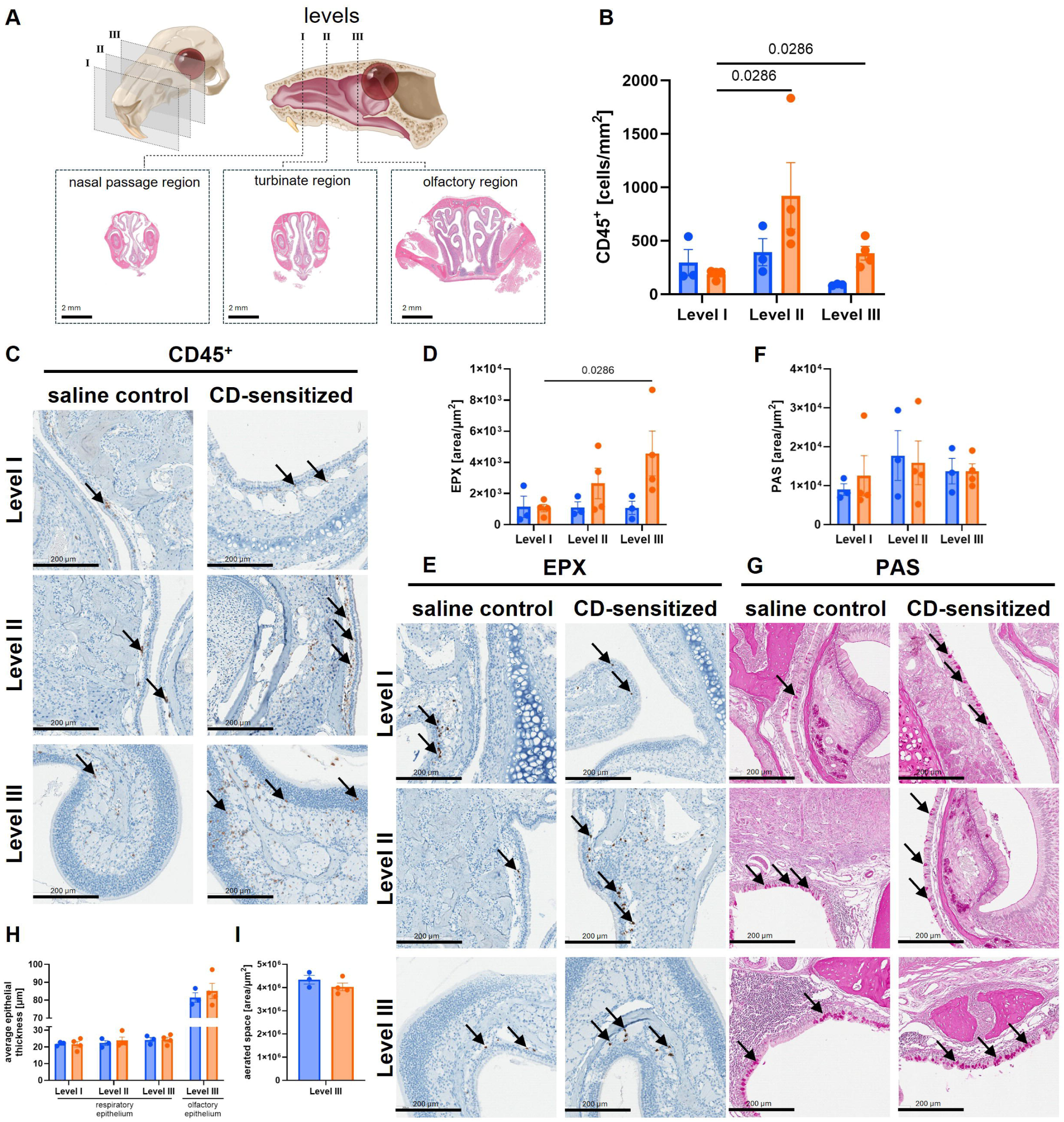
Most immunological changes in allergic rhinitis occur in nasal levels II and III. **(A)** Frontal skulls at 3h post-challenge were sectioned at Level 1 (nasal passage), Level II (naso- and maxillo-turbinates), and Level III (olfactory portion of the nasal cavity). **(B)** Quantification and **(C)** representative microscopic images of CD45^+^ immune cells in the nose. **(D)** Quantification and **(E)** representative microscopic images of eosinophil peroxidase (EPX) in the nose. **(F)** Quantification and **(G)** representative microscopic images of Period acid-Schiff (PAS) stained goblet cells. **(H)** Measurement of epithelial thickness. **(I)** Quantification of aerated space. Arrows demonstrate the presence of immuno-positive cells respectively granules (C, E) or PAS-positive goblet cells (G). n=3-4/group. Images are 20x magnification. Mann-Whitney tests.

Eosinophil peroxidase (EPX) is a cytotoxic protein specifically found in the granules of eosinophils (31). EPX expression was increased in CD-sensitized mice, with significantly higher presence in the olfactory region than in the nasal passage (**Fig 6D-E, S6**).

Mucin-producing goblet cells, as detected by Periodic acid Schiff (PAS) staining, showed similar numbers throughout all nasal levels in all mice (**Fig 6F-G**). Thickness of the respiratory epithelia in nasal passage and turbinate regions were similar between CD-sensitized and saline-control mice (**Fig 6H**). We assessed respiratory and olfactory epithelium thickness in Level III and observed minimally increased olfactory epithelial thickness in CD-sensitized mice that corresponded to decreased aerated space, albeit not with statistical significance (**Fig 6H-I**).

### The mouse model mimics allergic rhinitis features from the human clinical model

Direct comparison of this novel mouse model of cat allergic rhinitis with the human clinical model of experimental cat allergen exposure confirms analogous methodology and outcomes (**Table S6**): Sensitization in both humans and the experimentally exposed mice occurred through the nasal route. CD-sensitized mice and cat-allergic participants displayed significant increases in nasal symptoms following experimental exposure to allergen-diverse CD. Systemic sensitization was confirmed in both CD-sensitized mice and cat-allergic participants through increased serum total IgE. Systemic sensitization was further supported through an intradermal ear challenge in mice and skin testing in humans.

Exposure to CD resulted in increased nasal eosinophils in both CD-sensitized mice and cat-allergic participants. In mice, we were able to determine the localization of immune cells in the nasal cavity. CD-sensitized mice had higher numbers of CD45^+^ immune cells as well as increased EPX expression in the more caudal regions of the nose.

## DISCUSSION

To our knowledge, this is the first preclinical model of cat-AR directly informed by a human experimental system. The human model utilized the SPaC-EEU, a validated controlled allergen exposure facility for perennial allergens (23, 32), in which airborne CD is distributed and maintained in air at defined concentrations. Exposure levels in the SPaC-EEU (∼69ng Feld1/m³) fall within the range reported in other clinical models (25, 26) and are consistent with real-world airborne allergen levels in homes and public environments (26, 33) (**Table S2, Figure S7**).

Based on the human exposure data, we established a murine model using the same allergen source. Mice received 54ng of Feld1, corresponding to the inhaled dose during a 3-hour human exposure. The intranasal exposure volume (9µL) ensured localization to the upper airways (34, 35). While differences remain between airborne inhalation in humans and bolus intranasal administration in mice, this strategy provided a controlled and reproducible mucosal allergen exposure. Airborne exposures in chambers or with nose cones have been used in murine allergy and infection studies (36–38), however the sticky nature of allergens may limit consistent exposures with accurate assessment of inhaled allergen concentrations (39).

Following a 2-week intermittent sensitization and challenge schedule, mice demonstrated both local and systemic features of allergic sensitization. CD-sensitized mice displayed increased serum total IgE levels following IN challenge, consistent with systemic immune activation. A similar increase in serum total IgE was observed in cat-allergic participants following SPaC-EEU exposure, supporting cross-species alignment in systemic responses. Systemic responses were confirmed through an ear challenge assay, analogous to a human skin prick test, in which CD-sensitized mice exhibited prolonged ear swelling selectively in response to allergen injection. This response was absent in control mice and paralleled increased skin prick test reactivity in cat-allergic participants (23).

CD-sensitized mice exhibited increased nasal rubbing, replicating nasal itching in human participants following allergen exposure. While this finding supports functional relevance of the model, it is important to note that intranasal administration itself may contribute to nonspecific irritation. Indeed, both saline-control and CD-exposed mice showed early increases in nasal neutrophils and eosinophils compared to untreated controls, suggesting that procedural factors trigger initial immune responses. However, the increase in eosinophils and nasal rubbing observed at later time points specifically in CD-sensitized mice represent an allergen-driven late-phase immunological and clinical response.

To enable detailed analysis of mucosal immune responses, we developed a nasal sampling protocol that allows for efficient and reproducible collection of immune cells from the murine nasal cavity. In comparison to previously described methods that flush through the trachea or use smaller lavage volumes (40–43), our approach yielded substantially higher and more consistent cell numbers, improving the reliability of downstream analyses. We did not employ intravenous injection of αCD45 antibodies for intravascular versus tissue immune cell staining (44), because of expected epithelial cell damage, tight junction impairment, and vascular leakage in allergic inflammation, which could compromise staining localization.

Using the nasal cavity flushing and flow cytometry, dynamic changes in nasal immune cell composition following allergen exposure were assessed. Both saline-control and CD-sensitized mice showed an increase in the percentage of nasal neutrophils at 3h post challenge, consistent with an initial irritant response. Subsequently, only CD-sensitized mice demonstrated a late-phase increase in both percentage and counts of nasal eosinophils, replicating clinical observations of sustained eosinophilia following allergen challenge (45). Eosinophils are central effectors of type-2 inflammation (46) and are recruited to sites of allergic inflammation through epithelial-derived chemokines (47). At the site of inflammation, eosinophils release cytotoxic mediators including eosinophil cationic protein and EPX(31). While diurnal changes in eosinophil activity have been described (48), our sensitizations and challenges were all performed in the morning, hence we expect no diverging intrinsic impacts in our results. Nasal eosinophils did not increase at 3h post challenge in CD-sensitized mice. We suspect that the 3-hour timepoint captured a transition period in eosinophil function: eosinophils were activated in the early phase of the allergic response, promoted local inflammation and hence clinical signs (49, 50), and were then depleted. Concurrently, level-specific histological analyses captured increased eosinophil-specific EPX staining in CD-sensitized mice at 3h post challenge. CD11b and MHC II expression suggested a time-dependent recruitment and activation of eosinophils, with increased frequencies at later time points. These findings support the presence of a coordinated allergic response in the nasal mucosa.

Histological analysis revealed region-specific changes in the nasal cavity, including increased immune cell presence in turbinate and olfactory regions and elevated EPX levels in Level III. While increases in epithelial thickness and reductions in aerated space were observed, these changes did not reach statistical significance, suggesting that structural remodeling may require longer or repeated exposures. Nonetheless, these findings indicate that the model captures early features of tissue-level responses associated with allergic inflammation.

## CONCLUSION

We presented a clinically informed murine model of cat-AR that integrates natural allergen exposure, avoids the use of adjuvants, and captures key local and systemic features of allergic sensitization. It provides a robust and reproducible platform for mechanistic studies and therapeutic evaluation. By enabling investigation of cellular and molecular pathways in a physiologically relevant context, the here presented novel mouse model has the potential to advance our understanding of allergic rhinitis and support the development of improved therapeutic strategies.

## Supporting information

Car-AR Manuscript Supplement

## RESEARCH IMPACT

Based on an experimental human exposure model, we here developed a clinically informed mouse model of cat allergic rhinitis that recapitulates natural cat allergen exposure and core clinical and biological features of the human disease. This model will support investigations into the mechanisms of upper and associated lower respiratory tract allergic sensitization, and provide the basis for development of novel therapeutic approaches.

## AUTHOR CONTRIBUTIONS

Conceptualization: LH, CH, AKE, EK

Methodology: LH, WT, STN, JT, CH, AL, SH, VJ, TR, HB, LS, AJP, FC, TW, JSch, AKE, EK

Investigation: LH, WT, STN, JT, JSch, AKE, EK

Visualization: LH, WT, JT, AJP, FC, JSch, EK

Funding acquisition: AKE, EK

Project administration: STN, SG, CH, FC, JS, TW, LS, JSch, AKE, EK

Supervision: AKE, EK

Writing – original draft: LH, WT, EK

Writing – review & editing: LH, WT, JS, JSch, AKE, EK

## ACKNOWLEDGEMENTS

We sincerely thank the Kingston Allergy Research team for their support in implementing the clinical study, as well as the research participants for their invaluable contributions. We are grateful to the Queen’s University and RI-MUHC Animal Care Services teams for dedicated animal care, and to Jeff Mewburn at the DBMS Flow Cytometry Core Facility for expert technical support. We acknowledge Shakeel Virk and Nick Zhao (Queen’s University Department of Pathology and Molecular Medicine) for their assistance with HALO^®^ Link. Finally, we thank Dr. James Martin and Dr. Irah King (both McGill University) for their critical feedback on the manuscript, and we acknowledge our funders for their financial support of this work.

## DATA AVAILABILITY STATEMENT

All data and protocols are available upon request to the corresponding authors. The nasal rubbing code is available here: https://zenodo.org/records/19257529.

## ETHICAL APPROVAL STATEMENT

All clinical procedures were approved by the Queen’s University Health Sciences and Affiliated Teaching Hospitals Research Ethics Board. All participants provided informed consent. All animal studies were approved by the Animal Research Ethics Council at Queen’s University (protocol number: 2338) and McGill University (protocol number: MUHC-10263).

## REFERENCES

1. McCrory D, Williams J, Dolor R, Gray R, Kolimaga J, Reed S, et al. Management of allergic rhinitis in the working-age population. Evid Rep Technol Assess (Summ). 2003;67.

2. Sibbald; B, Rink; E. Epidemiology of seasonal and perennial rhinitis: clinical presentation and medical history. Thorax. 1991;46(12).

3. Salo P, Arbes S, Jaramillo R, Calatroni A, Weir C, Sever M, et al. Prevalence of allergic sensitization in the United States: results from the National Health and Nutrition Examination Survey (NHANES) 2005-2006. The Journal of allergy and clinical immunology. 2014;134(2).

4. Satyaraj E, Wedner HJ, Bousquet J. Keep the cat, change the care pathway: A transformational approach to managing Fel d 1, the major cat allergen. Allergy. 2019;74 Suppl 107(Suppl 107):5–17.

5. Bonnet B, Messaoudi K, Jacomet F, Michaud E, Fauquert JL, Caillaud D, et al. An update on molecular cat allergens: Fel d 1 and what else? Chapter 1: Fel d 1, the major cat allergen. Allergy Asthma Clin Immunol. 2018;14(1):14.

6. De Lucca SD, O’Meara T J, Tovey ER. Exposure to mite and cat allergens on a range of clothing items at home and the transfer of cat allergen in the workplace. J Allergy Clin Immunol. 2000;106(5):874–9.

7. Small P, Keith PK, Kim H. Allergic rhinitis. Allergy Asthma Clin Immunol. 2018;14(Suppl 2):51.

8. Meltzer EO, Bukstein DA. The economic impact of allergic rhinitis and current guidelines for treatment. Ann Allergy Asthma Immunol. 2011;106(2 Suppl):S12–6.

9. Arbes; SJ, Gergen; PJ, Vaughn; B, Zeldin; DC. Asthma Cases Attributable to Atopy: Results from the Third National Health and Nutrition Examination Survey. The Journal of allergy and clinical immunology. 2007;120(5).

10. Noertjojo; K, Dimich-Ward; H, Obata; H, Manfreda; J, Chan-Yeung; M. Exposure and sensitization to cat dander: Asthma and asthma-like symptoms among adults. Journal of Allergy and Clinical Immunology. 1999;103(1).

11. Mudarri DH. Valuing the Economic Costs of Allergic Rhinitis, Acute Bronchitis, and Asthma from Exposure to Indoor Dampness and Mold in the US. J Environ Public Health. 2016;2016:2386596.

12. Wood R, Chapman M, Adkinson N, Eggleston P. The effect of cat removal on allergen content in household-dust samples - PubMed. The Journal of Allergy and Clinical Immunology. 1989;83(4).

13. Arbes SJ, Cohn RD, Yin M, Muilenberg ML, Friedman W, Zeldin DC. Dog allergen (Can f 1) and cat allergen (Fel d 1) in US homes: Results from the National Survey of Lead and Allergens in Housing. Annals of allergy, asthma & immunology. 2008;101(5).

14. Niesler A, Scigala G, Ludzen-Izbinska B. Cat (Fel d 1) and dog (Can f 1) allergen levels in cars, dwellings and schools. Aerobiologia (Bologna). 2016;32(3):571–80.

15. Nelson M, Cox L. Chapter 9: Allergen Immunotherapy Extract Preparation Manual. AAAAI Practice Management Resource Guide. 2014.

16. Van Metre TE, Jr., Marsh DG, Adkinson NF, Jr., Kagey-Sobotka A, Khattignavong A, Norman PS, Jr., et al. Immunotherapy decreases skin sensitivity to cat extract. J Allergy Clin Immunol. 1989;83(5):888–99.

17. Jimenez-Blanco MA, Gonzalez-Mendiola MR, Boteanu C, Elera JD, Sanchez-Millan ML, Ruiz-Garcia M, et al. Effectiveness and safety of subcutaneous immunotherapy using a depigmented, polymerized extract of cat epithelium in allergic patients: a retrospective, real-world study. Front Allergy. 2025;6:1642315.

18. van Hage M, Kack U, Asarnoj A, Konradsen JR. An update on the prevalence and diagnosis of cat and dog allergy -Emphasizing the role of molecular allergy diagnostics. Mol Immunol. 2023;157:1–7.

19. Layhadi JA, Gutierrez LC, Keane ST, Fulton W, Samson NA, Wu LYD, et al. Fel d 1-Expressing Plant-Derived Bioparticle: A Novel Treatment for Cat Allergy. Allergy. 2026.

20. Wu L, Aglas L, Van Ree R, Tropper G, Gutierrez BC, Shamji M, et al. Fel D 1 Plant Enveloped Bioparticles, a Candidate Cat Allergy Immunotherapy with Excellent Hypoallergenic Properties. Annals of Allergy, Asthma & Immunology. 2024;133(6):S21–S2.

21. Busold S, Aglas L, Menage C, Auger L, Desgagnes R, Faye L, et al. Fel d 1 surface expression on plant-made eBioparticles combines potent immune activation and hypoallergenicity. Allergy. 2022;77(10):3124–6.

22. Jones JT, Tassew DD, Herrera LK, Walton-Filipczak SR, Montera MA, Chand HS, et al. Extent of allergic inflammation depends on intermittent versus continuous sensitization to house dust mite. Inhal Toxicol. 2017;29(3):106–12.

23. Hossenbaccus L, Linton S, Garvey S, Botting H, Walker T, Steacy L, et al. Clinical validation of controlled exposure to cat dander in the Specialized Particulate Control Environmental Exposure Unit (SPaC-EEU). Allergy Asthma Clin Immunol. 2025;21(1):33.

24. Bankhead P, Loughrey MB, Fernandez JA, Dombrowski Y, McArt DG, Dunne PD, et al. QuPath: Open source software for digital pathology image analysis. Sci Rep. 2017;7(1):16878.

25. Wood RA, Laheri AN, Eggleston PA. The aerodynamic characteristics of cat allergen. Clin Exp Allergy. 1993;23(9):733–9.

26. Bollinger ME, Eggleston PA, Flanagan E, Wood RA. Cat antigen in homes with and without cats may induce allergic symptoms. J Allergy Clin Immunol. 1996;97(4):907–14.

27. Pulmonary Anatomy and Physiology: The Basics. Principles of Pulmonary Medicine. 2014.

28. Jia GQ, Gonzalo JA, Hidalgo A, Wagner D, Cybulsky M, Gutierrez-Ramos JC. Selective eosinophil transendothelial migration triggered by eotaxin via modulation of Mac-1/ICAM-1 and VLA-4/VCAM-1 interactions. Int Immunol. 1999;11(1):1–10.

29. Roche PA, Furuta K. The ins and outs of MHC class II-mediated antigen processing and presentation. Nat Rev Immunol. 2015;15(4):203–16.

30. National Institute of Environmental Health Sciences. Nasal Cavity: Revised Guides for Organ Sampling and Trimming in Rats and Mice.

31. Tang M, Charbit AR, Johansson MW, Jarjour NN, Denlinger LC, Raymond WW, et al. Utility of eosinophil peroxidase as a biomarker of eosinophilic inflammation in asthma. J Allergy Clin Immunol. 2024;154(3):580–91 e6.

32. Hossenbaccus L, Linton S, Thiele J, Steacy L, Walker T, Malone C, et al. Clinical validation of controlled exposure to house dust mite in the environmental exposure unit (EEU). Allergy Asthma Clin Immunol. 2021;17(1):34.

33. Custovic A, Fletcher A, Pickering CA, Francis HC, Green R, Smith A, et al. Domestic allergens in public places III: house dust mite, cat, dog and cockroach allergens in British hospitals. Clin Exp Allergy. 1998;28(1):53–9.

34. McCusker C, Chicoine M, Hamid Q, Mazer B. Site-specific sensitization in a murine model of allergic rhinitis: role of the upper airway in lower airways disease. J Allergy Clin Immunol. 2002;110(6):891–8.

35. Shibamori M, Ogino K, Kambayashi Y, Ishiyama H. Intranasal mite allergen induces allergic asthma-like responses in NC/Nga mice. Life Sci. 2006;78(9):987–94.

36. Templeton SP, Buskirk AD, Green BJ, Beezhold DH, Schmechel D. Murine models of airway fungal exposure and allergic sensitization. Med Mycol. 2010;48(2):217–28.

37. Belser JA, Gustin KM, Katz JM, Maines TR, Tumpey TM. Comparison of traditional intranasal and aerosol inhalation inoculation of mice with influenza A viruses. Virology. 2015;481:107–12.

38. Lucci F, Tan WT, Krishnan S, Hoeng J, Vanscheeuwijck P, Jaeger R, et al. Experimental and computational investigation of a nose-only exposure chamber. Aerosol Science and Technology. 2019;54(3):277–90.

39. Weber RW. The Nature of Allergens. Immunology and Allergy Clinics of North America. 1987;7(2):191– 203.

40. Schenck LP, McGrath JJC, Lamarche D, Stampfli MR, Bowdish DME, Surette MG. Nasal Tissue Extraction Is Essential for Characterization of the Murine Upper Respiratory Tract Microbiota. mSphere. 2020;5(6).

41. Puchta A, Verschoor CP, Thurn T, Bowdish DM. Characterization of inflammatory responses during intranasal colonization with Streptococcus pneumoniae. J Vis Exp. 2014(83):e50490.

42. Hernani Mde L, Ferreira PC, Ferreira DM, Miyaji EN, Ho PL, Oliveira ML. Nasal immunization of mice with Lactobacillus casei expressing the pneumococcal surface protein C primes the immune system and decreases pneumococcal nasopharyngeal colonization in mice. FEMS Immunol Med Microbiol. 2011;62(3):263–72.

43. McInnes EF, Rasmussen L, Fung P, Auld AM, Alvarez L, Lawrence DA, et al. Prevalence of viral, bacterial and parasitological diseases in rats and mice used in research environments in Australasia over a 5-y period. Lab Anim (NY). 2011;40(11):341–50.

44. Anderson KG, Mayer-Barber K, Sung H, Beura L, James BR, Taylor JJ, et al. Intravascular staining for discrimination of vascular and tissue leukocytes. Nat Protoc. 2014;9(1):209–22.

45. Terada N, Hamano N, Kim WJ, Hirai K, Nakajima T, Yamada H, et al. The kinetics of allergen-induced eotaxin level in nasal lavage fluid: its key role in eosinophil recruitment in nasal mucosa. Am J Respir Crit Care Med. 2001;164(4):575–9.

46. Matsunaga K, Koarai A, Koto H, Shirai T, Muraki M, Yamaguchi M, et al. Guidance for type 2 inflammatory biomarkers. Respir Investig. 2025;63(3):273–88.

47. Zajkowska M, Mroczko B. From Allergy to Cancer-Clinical Usefulness of Eotaxins. Cancers (Basel). 2021;13(1).

48. Baumann A, Gonnenwein S, Bischoff SC, Sherman H, Chapnik N, Froy O, et al. The circadian clock is functional in eosinophils and mast cells. Immunology. 2013;140(4):465–74.

49. Clark K, Simson L, Newcombe N, Koskinen AM, Mattes J, Lee NA, et al. Eosinophil degranulation in the allergic lung of mice primarily occurs in the airway lumen. J Leukoc Biol. 2004;75(6):1001–9.

50. Hamour AF, Lee JJ, Wasilewski E, Monteiro E, Lee JM, Vescan A, et al. Murine model for chronic rhinosinusitis: an interventional study. J Otolaryngol Head Neck Surg. 2023;52(1):32.

