## Supplementary material for "Development and translational validation of a novel mouse model for allergic rhinitis": Car-AR Manuscript Supplement

**SUPPLEMENTAL MATERIALS**

- Supplemental methods
- Table S1. Overview of murine models of cat-AR.
- Table S2. Fel d 1 concentrations in air samples from experimental and natural exposures.
- Table S3. Time chart of clinical study procedures.
- Table S4. Scoring of nasal symptoms in clinical study.
- Table S5. Criteria for appearance and activity scoring for health monitoring of mice.
- Table S6. Comparison of human and mouse models.
- Fig S1. Nasal symptoms after human experimental cat dander exposure.
- Fig S2. Eosinophil detection in human nasal lavage samples.
- Fig S3. Example of nasal rubbing in mice.
- Fig S4. Total numbers of nasal immune cells in CD-sensitized mice.
- Fig S5. Flowcytometric detection of CD11b and MHC II expression on murine nasal eosinophils.
- Fig S6. Nasal localization of EPX in mice.
- Fig S7. Visual representation of Fel d 1 exposure concentrations.
- Fig S8. Measurement of epithelial thickness in the murine nasal mucosa.
- Fig S9. Annotation of the aerated area at Level III of the mouse nose.
- Supplemental references
- GitHub Repository README

**Supplemental material for:**

### Development and translational validation of a novel mouse model for cat allergic rhinitis

Lubnaa Hossenbaccus, Wil P. Taylor, Sara Teimouri Nezhad, Jack M. Taylor, Cortney Haird, Andisheh Liaghat, Sarah E. Hopkins, Vidthiya Jeyanathan, Tyson Rudolf, Adrian J. Paz, Hannah Botting, Lisa M. Steacy, Fazila Chouiali, Terry Walker, Joaquin Sanz, Sarah Garvey, Jan Schinköthe, Anne K. Ellis, Eva Kaufmann

#### Supplemental Methods: Methodological details

##### Study design

Human experimental cat allergen exposure was used to define the temporal trajectory of clinical symptoms and upper airway immune responses in allergic patients. Cat-allergic and non-allergic participants were challenged with airborne cat dander allergen in the Specialized Particulate Control Environmental Exposure Unit (SPaC-EEU), a validated inhalational exposure facility for inducing allergic rhinitis (1-3). Nasal symptom scores were recorded at predefined intervals pre- and post-challenge, and biospecimens were collected for serum IgE (IgE) and nasal immune cell profiling through ImmunoCAP analysis and manual cell counting, respectively. Guided by these kinetics, we developed a mouse model of cat allergen induced allergic rhinitis using an allergen dose calibrated to human exposure (**Table S2**) which was administered strictly intranasally in intermitted intervals. Ultimately, mice underwent a controlled intranasal challenge, with serial sampling to quantify nasal immune-cell dynamics and symptomatic outcomes. Concordance between species was assessed by aligning time-resolved cellular and symptom endpoints and by cross-modal analyses of shared signatures. This preclinical platform enables mechanistic interrogation and therapeutic testing in insightful ways not feasible in clinical research. We further provide a code for simplified analysis of nasal rubbing as clinical sign of allergic rhinitis in mice. Our data highlight the importance of per level investigation of nasal immune changes.

**Cat dander allergen**

Cat dander was obtained from Stallergenes Greer, USA (item number: RME63P; lot number: 409233-1). For human experimental exposures, the allergen was lofted into the air of the SPaC-EEU. For mouse experiments, the cat dander powder was reconstituted with Dulbecco's Phosphate-Buffered Saline (Life Technologies) containing 0.05% Tween-20 (DPBS-T) (MP Biomedicals LLC) and was stored at 2ºC. The Fel d 1 concentration in the reconstituted cat dander preparation was determined by a Fel d 1-specific ELISA (Indoor Biotechnologies).

**Human clinical study**

All procedures were approved by Queen's University Health Sciences and Affiliated Teaching Hospitals Research Ethics Board. All participants provided informed consent. Detailed protocols for the human cat dander exposure study, including but not limited to, screening procedures, inclusion/exclusion criteria, and medication washout periods, have been previously published (2). Briefly, 31 confirmed cat-allergic and 15 non-allergic participants between the ages of 12 and 65 years were included in this study. Participants with a minimum 2-year clinical history of allergy symptoms to cats and a positive skin prick test to cat hair were enrolled as "cat-allergic" and those with no history of cat allergies and a negative skin prick test to all tested allergens (cat hair, birch, timothy grass, short ragweed, *D. pteronyssinus*, *D. farinae*, dog epithelium, and mold mix) were enrolled as "non-allergic". All participants were required to be able and willing to provide written informed consent or assent and to comply with study requirements. Participants were exposed to airborne cat dander in the SPaC-EEU for 3 hours in one of two exposure sessions, due to space limitations. Pre- and post-exposure symptom scores and biological samples were collected at various timepoints (**Table S3**).

Specialized Particulate Controlled Environmental Exposure Unit (SPaC-EEU)

The SPaC-EEU is a specifically designed perennial allergen exposure facility designed within the larger Environmental Exposure Unit (EEU) in the Kingson Health Science Centre. The setup and methodology of the EEU have informed standardization guidelines for allergen exposure facilities published by the European Academy of Allergy and Clinical Immunology (EAACI) (4, 5). The SPaC-EEU has been validated for use with house dust mite exposure and cat allergen exposure (1-3, 6, 7). For the here presented study, air samples were collected from air sampling cassettes at the front, middle, and rear of the facility, revealing a mean exposure Fel d 1 concentration of 69 $ng/m^3$.

Symptom scoring

All participants were trained to assess nasal symptoms on a scale from 0 to 3, increasing in severity (**Table S4**). No partial scores were permitted. The symptom scores were recorded on paper diary cards pre-exposure (baseline), at 15 minutes, at 30 minutes, then every half-hour throughout the exposure, and then on an hourly basis up to 12 hours, and again at 24 hours after the onset of allergen exposure (**Table S3**). Symptoms of sneezing, runny nose/post-nasal drip, nasal congestion/stuffiness, and itchy nose were tallied as Total Nasal Symptom Score out of 12.

Blood collection, serum isolation, and IgE evaluation

Human blood samples were collected through venipuncture in 3.5 mL SST tubes (Becton Dickinson) from both cat-allergic and non-allergic participants pre- (baseline) and post-exposure (3-, 6-, and 24-hours) (**Table S3**). Blood was allowed to clot for 30-60 minutes at room temperature and then the SST tubes were centrifuged at 1,500 g for 15 minutes at RT. The serum was aliquoted into microcentrifuge tubes (Eppendorf) and stored at -80°C.

Serum samples were used for specific and total IgE analysis using an ImmunoCAP assay (Thermo Fisher Scientific) run on the Phadia™ 200 instrument, with all assay procedures completed by a clinically validated automation instrument.

<u>Nasal lavage collection, sample processing, and cell counting</u>

Nasal lavage samples were collected from a randomly determined subset of 10 cat-allergic and 4 non-allergic participants pre- (baseline) and post-exposure (3- and 24-hours), as previously described (8, 9). Briefly, nasal lavage samples were collected by a trained staff member. An appropriately sized sterile nasal olive attached to a syringe filled with 5.0 mL of sterile saline was inserted into one nostril, ensuring a sealed fit. The saline was gently flushed in and out of the same nostril approximately 20 times, back and forth into the same syringe. The collected nasal fluid was put into a pre-weighed 15-mL polystyrene tube and placed on ice.

Tubes were re-weighed to determine sample volume; 1% dithiothreitol (1:10 w/v; Sigma-Aldrich) was subsequently added. Tubes were rocked on a shaker for 15 minutes at RT, followed by centrifugation at 600 g for 4 minutes at RT. Excess supernatant, except for ~1 mL was discarded and the cellular pellet was resuspended in the remaining 1mL. Cells were counted and samples were diluted with DPBS to achieve a concentration of 200,000 cells/mL. Microscope slides were pre-wet with 50 µL DPBS, spun at 300 rpm for 3 minutes using Shandon Cytospin 4 (Thermo Fisher Scientific). Cytospin funnels were then loaded with 100 µL of samples, and samples were spun onto pre-wet slides at 300 rpm for 5 minutes. Slides were airdried for 24 hours.

Cytospin slides were stained with the Hemacolor® Stain Set (MilliporeSigma, product number: 65044-M) and left to dry overnight, prior to mounting with Permount™ Mounting Medium (Fisher

Scientific).

Samples on slides were scanned using the Aperio VERSA Scanner (Leica) at 20x or 40x magnification with brightfield settings. Digital renderings of slides were uploaded onto HALO (Indica Labs), courtesy of the Queen's Laboratory for Molecular Pathology. Slides were manually counted for neutrophils, eosinophils, monocytes, lymphocytes, basophils, and epithelial cells. Up to 400 cells per slide were counted by 5 trained cell counters on HALO Link (Indica Labs), all blinded to allergic status. Quality assurance of identified cells was completed, and cell numbers were analyzed as percentage of counted cells and as a percentage of immune cells.

#### Mouse experiments

##### Mice

Male and female 7- to 12-week-old BALB/c and C57BL/6 mice were purchased from Jackson Laboratories or bred at Queen's University Animal Facility, or the Animal Care Service Facility of the Research Institute of the McGill University Health Centre (RI-MUHC). Mice were housed under specific pathogen-free conditions. All animal studies were approved by the Animal Research Ethics Council at Queen's University (protocol number: 2338) and McGill University (protocol number: MUHC-10263).

##### Mouse model of cat allergic rhinitis

The human clinical exposures in the SPaC-EEU had mean Fel d 1 concentrations of 69 ng/m$^3$. We calculated that exposure to rounded 50 ng/m$^3$ over a 3-hour exposure period led to participants inhaling approximately 54 ng of Fel d 1:

$$\begin{pmatrix} average\ human \\ inhalation\ rate \\ as\ L/minute \end{pmatrix}\left(\frac{60\ min}{hr}\right)\left(\frac{m^3}{1000L}\right)\left(\frac{ng}{m^3}\right)\begin{pmatrix} total\ exposure\ time \\ in\ hours \end{pmatrix} = ng\ of\ Fel\ d\ 1$$

$$\left(\frac{6\ L}{min}\right)\left(\frac{60\ min}{hr}\right)\left(\frac{m^3}{1000L}\right)\left(\frac{50\ ng}{m^3}\right)(3\ hours) = 54\ ng\ of\ Fel\ d\ 1$$

As mice are more tolerant to immunological stimuli compared to humans (10), and often human-equivalent doses are used for mice (11-14), cat dander (CD)-sensitized mice were intranasally (IN) exposed to 54 ng of Fel d 1 (experiment days 1, 2, 3; 7, 8, and 9) in 9 µL of reconstituted cat dander under isoflurane anesthesia. "No treatment" (NT) control mice underwent the same isoflurane anesthesia with no IN exposure, whereas saline control mice received isoflurane anesthesia and IN sterile PBS (Wisent) at the same intervals. On day 13, CD-sensitized and saline control mice received an IN challenge with their respective stimuli. All mouse experiments were performed in the morning.

Weights were collected before each IN administration. Mice were video recorded for a 30-minute period after each sensitization to observe behavior and record instances of nasal rubbing.

Mice were euthanized by $CO_2$ asphyxiation under deep isoflurane anesthesia. Tissue samples were collected from CD-sensitized and saline control mice at 3- and 6-hours post-challenge, with sex- and age-matched no treatment control mice as baseline.

<u>Health monitoring</u>

Mice were monitored for 30 minutes after each sensitization and challenge period for changes in their health and activity due to exposures. Appearance scores (/3) were based on posture, coat, and eye appearance. Activity scores (/3) were based on alertness and responsiveness (**Table S5**).

Distribution of fluid following intranasal administration

To delineate the anatomical extent of fluid inhalation in IN allergen exposure, 9 μL of Evan's Blue solution (0.125% w/v in PBS (15)) was administered to isoflurane-anesthetized naïve mice. After 5 minutes and once fully awakened, mice were anesthetized with isoflurane again and directly euthanized by $CO_2$ asphyxiation.

The head and chest of the mouse were sprayed with 70% ethanol, and the mandible was removed as described in *Nasal Fluid Collection Method.* To visualize the nasal cavity, the hard and soft palate were removed by making two parallel incisions from the nasopharynx toward the nose as close as possible to the buccal side of the upper molars. Using scissors, an incision was made through the skin, from the level of the trachea until the xiphoid process of the sternum. The skin was opened laterally on both sides to reveal the underlying muscles of the neck and thorax. The sternocleidomastoids, hyoid muscles, and remaining soft tissue were removed using micro dissecting scissors to reveal the trachea and esophagus. To display the lung *in situ*, a cut was made through the abdomen along the diaphragmatic border to approximately the mid axillary line. The diaphragm was removed. A vertical cut was made on each side of the sternum from the xiphoid process to the manubriosternal joint. The two sides of the ribcage were opened laterally to reveal the lung *in situ*. The heart was removed to visualize the main bronchi. Trachea and lung were opened through medial horizontal dissection. Evan's Blue localization was documented photographically.

Intradermal ear challenge

A subset of CD-sensitized and saline control mice received an intradermal ear challenge, instead of intranasal challenge on day 13. To that end, mice were anesthetized with isoflurane and sterile PBS was administered intradermally in the left ear using a 31G needle, and cat dander allergen, with 54 ng of Fel d 1, was injected intradermally in the right ear. Ear thickness measurements were collected using a digital micrometer (Beslands, accuracy 0.003 mm) prior to injection challenge (baseline), immediately after injection challenge (t=0), hourly for up to 12 hours, and at 24 hours post-challenge.

Mouse serum isolation

Blood samples were collected from anesthetized mice via cardiac puncture into lithium heparin tubes (BD, catalog number: 13-680-62). Blood samples were allowed to clot for 45 minutes at room temperature. Tubes were then centrifuged for 90 seconds at 15,000g. Serum was pipetted into labelled Eppendorf tubes and stored at -20°C until further processing.

Mouse serum IgE quantification

The ELISA MAX™ Standard Set Mouse IgE (Biolegend) was used to determine concentrations of total IgE in mouse serum samples (diluted 1:40 and/or 1:50), according to the manufacturer's instructions. Attempts to modify the total IgE ELISA for CD- or Fel d 1 IgE specificity were unsuccessful.

Nasal fluid collection method

Following $CO_2$ euthanasia, mice were sprayed with 70% ethanol. An incision was made beside the ear, and the skin was cut around the base of the neck, and to the base of the shoulder. Using sharp scissors, the head and neck of the mouse was severed at the level of T1. The mandible was removed by inserting the scissors into the mouth and cutting horizontally through the cheeks parallel to the jaw line toward the neck. Excess tissue was removed until the white triangle of the soft palate was visible to indicate the position of the nasopharyngeal sphincter at the apex of the palate. A 22G feeding tube (Instech Laboratories) connected to a 10 mL syringe filled with 10 mL of sterile saline (Wisent) was inserted into the nasopharyngeal sphincter. With the mouse head held vertically, nose down, above a 50 mL conical tube, the plunger was slowly depressed to flush the nasal cavity, avoiding spillover that would rinse the oral side of the soft and hard palate. This technique led to the flushing of both nares simultaneously. The flushing of the nasal cavity was repeated with an additional 10 mL of sterile saline using a 20G feeding tube (Instech Laboratories). Both nasal flushing samples from each mouse were collected in the same conical tube.

Processing of nasal fluid samples

Nasal fluid samples were spun at 1600 rpm for 10 minutes. The supernatant was decanted. Cell pellets were resuspended in 1 mL RBC lysis buffer for 1 minute before the addition of 10 mL of RPMI (Wisent) to each tube. Samples were then transferred into a 15 mL tube and spun at 1600 rpm for 10 minutes. The supernatant was aspirated and the pellet was resuspended in 0.5 mL RPMI. Cells were counted using Trypan Blue in a 1:2 dilution.

Flow cytometry

Nasal cell preparations were transferred into v-bottom 96-well plates and stained with fixable viability stain eFluor501 (Invitrogen) at a dilution of 1:1000 for 30 minutes at 4°C. Cells were washed with sterile PBS (Wisent) supplemented with 0.5% BSA (Wisent) at 4°C for 6 minutes, then incubated with Fc block (clone: 93, Invitrogen) at 1:100 concentration for 10 minutes at 4°C.

Cells were next incubated with fluorescently labelled antibodies for 30 minutes at 4°C.

Innate panel antibodies included anti-c-kit-V450 (1:100; clone: 2B8, Biolegend), anti-CD11c-BV786 (1:100; clone: HL3, BD Horizon), anti-CD45.2-FITC (1:50; clone: 104, Biolegend), anti-Ly6G PerCP-Cy5.5 (1:100; clone:1A8, Biolegend), anti-CCR2-PE (1:100; clone: SA203G11, Biolegend), anti-Siglec-F-PE-CF594 (1:100; clone: E50-2440, BD Horizon), anti-MHC-II-BUV737 (1:100; Clone: M5/114.15.2, BD), anti-CD11b-PE-Cy7 (1:100; clone: M1/70, Biolegend), anti-Ly6C-APC (1:100; clone: AL-21, BD Horizon), anti-CD49b-BV650 (1:100; clone: HMa2, BD Horizon), anti-FcεRIa-AF700 (1:100; clone: MAR-1, Biolegend), anti-F4/80-APC-Cy7 (1:100; clone: BM8, Biolegend), and anti-IL5R-BUV395 (1:100; clone: T21, BD).

Adaptive panel antibodies included anti-CD4-V450 (1:100; CLONE; Biolegend), anti-CD45.2 FITC (1:50; clone: 104, Biolegend), anti-Siglec-F-PE (1:100; clone: S17007L, Biolegend), anti-γδ-BUV395 (1:100; clone: GL3, BD), anti-NK1.1-BV650 (1:100; clone: PK136, Invitrogen), anti-CD3-APC (1:100; clone: 145-2C11, Biolegend) anti-CD19-APC-Cy7 (1:100; clone: eBio1D3, Invitrogen), anti-CD11b-PE-Cy7 (1:100; clone: M1/70, Biolegend), anti-CD8-BV786 (1:100; clone: 53-6.7, BD), anti-CD62L-AF700 (1:100; clone: MEL-14, Biolegend), and anti-CD44-PE-CF594 (1:100; clone: IM7, BD).

Subsequently, all cells were washed with PBS/0.5% BSA and resuspended in 1% paraformaldehyde (Fisher). Cells were acquired on CytoFLEX (Beckman Coulter) or FACSymphony A5 flow cytometers (BD Biosciences). Analyses were performed using FlowJo software v.10.1 (Treestar).

Neutrophils were defined as live single cells, $CD45^+$, $SiglecF^-$, $CD11b^+$, $Ly6G^+$, $Ly6C^+$. Eosinophils: live single cells, $CD45^+$, $Ly6G^-$, $F4/80^{int/hi}$, $IL\text{-}5R^+$, $SigF^{int/hi}$. Within this population, expression levels for CD11b vs MHC II were quantified. Basophils were defined as live single cells, $CD45^+$, $MHCII^-$, $cKit^-$, $CD49b^+$, $Fc\varepsilon RI^+$. Mast cells: live single cells, $CD45^+$, $MHCII^-$, $cKit^+$, $CD11b^+$. T cells: live single cells, $CD45^+$, $CD3^+$, $CD19^-$. B cells: live single cells, $CD45^+$, $CD19^+$, $CD3^-$. NK cells: live single cells, $CD45^+$, $SiglecF^-$, $CD11b^-$, $NK1.1^+$. Macrophages: live single cells, $CD45^+$, $SiglecF^-$, $CD11b^+$, $Ly6G^-$, $Ly6C^{lo/int}$, $F4/80^{int/hi}$, $CD11c^-$.

Histology

*Processing*

Decapitation and removal of the mandible was performed following the same procedure as described for nasal fluid collection. The soft tissue of the head was stripped using forceps and blunt dissection. The eyes and optic nerves were removed from the orbits. Using a surgical scalpel, a sagittal cut was made through the skull at the lambdoid suture. The brain was removed with a probe through the sagittal opening. Tissue-stripped skulls were fixed in 4% formaldehyde for 24 hours, after which they were moved into 70% ethanol.

Skulls were then placed in a decalcifying agent (Fisher Scientific; catalog number 50-255-2444) for 16 hours. The skulls were trimmed through the removal of the anterior and posterior portions, using a razor blade, producing an ~1 cm length of nasal sinus. The rostral skull was sectioned at the Level 1 (nasal passages), Level II (naso- and maxillo-turbinates), and Level III (olfactory portion of the nasal cavity) (16). Formaldehyde-fixed, paraffin-embedded (FFPE) blocks were produced and sectioned at 5 μm thickness. Processing of FFPE-blocks and staining was performed by the Histopathology Core of RI-MUHC, Montreal, QC, Canada.

*Histological and immunohistochemical staining*

For routine histology, sections were stained with hematoxylin & eosin (Leica CV5030-ST5020) according to the manufacturer's instructions. For immunohistochemical analysis, staining was performed using the Discovery Ultra auto stainer (Roche). Serial sections for all three rostral skull levels were prepared; the first cut was stained with PAS (to quantify goblet cells), the second with Hematoxylin + EPX- 3,3'-diaminobenzidine (DAB) (to quantify eosinophils), and the third with Hematoxylin + CD45-DAB to quantify immune cells. To that end, sections of 4 μm thickness were deparaffinized and rehydrated. After antigen retrieval treatment in citrate buffer (Roche, pH 6.0, 32 min), sections were incubated for 32 minutes at 37°C with anti-mouse EPX (1:150, Cell Signalling Technology$^{R}$, catalogue number 98757, monoclonal) or anti-mouse CD45 (1:300, Cell Signalling Technology$^{R}$, catalogue number 70257, monoclonal). A secondary antibody incubation was completed with OmniMap anti-RB HRP (Roche, catalogue number 760-4311) for 20 minutes at RT, followed by detection ChromoMap DAB (Roche, catalogue number 760-159). Slides were then counterstained with Hematoxylin (760-2021, Roche) and cover slipped. Slides were digitally scanned until 40x (Leica Aperio Turbo) for morphometric analysis.

*Whole-slide image analysis*

Whole-slide images (WSIs) were analyzed by a board-certified veterinary pathologist (JSch, Dipl. ACVP) using QuPath (version 6.0.0) (17) who was blinded to the experimental groups at the time of analysis. For each staining, a separate project was established to permit batch quantification of all corresponding slides. Before quantitative analysis, the Estimate Stain Vectors function was applied to digitally unmix the chromogens, thereby distinguishing hematoxylin (blue) from DAB (brown) or PAS (pink-purple), as appropriate.

The respiratory and olfactory mucosa, including the adjacent submucosa, were defined as regions of interest. To generate these regions of interest, a pixel classifier was trained on manually annotated areas to discriminate "tissue" from "background." This classifier produced a mucosa/submucosa object for each nasal coronal section, which was subsequently manually refined to exclude the lumen, cartilage, bone, large glands, medium- to large-caliber vessels, and tissue artifacts (e.g., folds). The resulting refined annotations served as the basis for all subsequent measurements.

EPX immunoreactivity was quantified using a trained pixel classifier and expressed as the EPX-positive area fraction within the annotated region of interest. An analogous approach was applied to PAS-stained WSIs, with results reported as the PAS-positive area fraction within the region of interest.

For CD45 immunostaining, positively labeled cells were quantified using the Positive Cell Detection algorithm with the following parameters: hematoxylin optical density (OD) pixel size, 0.5 µm; nucleus background radius, 8 µm; sigma, 1.8 µm; minimum nucleus area, 10 $µm^2$; maximum nucleus area, 120 $µm^2$; cell expansion, 3 µm; and DAB OD mean threshold, 0.08. CD45 positive cells were normalized to tissue area and expressed as positive cells/$mm^2$.

Epithelial thickness in the respiratory and olfactory mucosa was measured on H&E-stained whole-slide images in a standardized manner, adapted from Sánchez-Montalvo et al. (18). At each nasal coronal level, six regions of interest were defined for the respiratory mucosa and six for the olfactory mucosa (level III only). Within each region of interest, epithelial thickness was measured at three sites by drawing a perpendicular line from the basement membrane to the apical cytoplasmic border (**Fig S8**), yielding 18 measurements per mucosal compartment and coronal level.

Aerated area was quantified at nasal coronal Level III using the Create Thresholder function. Two classes, "aerated area" and "tissue," were defined and separated using a single threshold of 229, with a minimum object size of 100,000 $\mu m^2$ and a minimum hole size of 1,000,000 $\mu m^2$. This approach generated a contiguous annotation of the aerated compartment, which was quantified as total aerated area in $\mu m^2$ (**Fig S9**).

<u>AI-assisted behavioural analysis of nasal rubbing in mice</u>

*Video acquisition*

Mice were video recorded in their home cages for 30 minutes following each exposure unless otherwise indicated. Four cages (one mouse per cage) were placed in a biological safety cabinet with open lids, and recording was performed using a fixed smartphone camera mounted on the cabinet glass. Cage positions were rotated between experiments to avoid positional bias. Videos were collected in M4V and totaled approximately 52 hours of footage.

*Behavioural event selection and annotation*

To enable objective and blinded behavioural quantification, we developed a containerized, local-first video analysis and annotation tool to identify candidate nasal rubbing events. Nasal rubbing was operationally defined as lifting of the forepaws toward the nose region, a movement associated with nasal irritation but also observed during grooming or feeding. The tool reduced video complexity by isolating mouse-specific regions of interest and generating cropped clips for annotation. All candidate events were manually reviewed and classified by a blinded researcher to distinguish nasal rubbing from grooming, feeding, or unrelated movements. An episode of nasal rubbing was characterized by the starting of nasal rubbing behaviour until termination, independent of the specific number of individual rubs on the nose.

*Software architecture and deployment*

The AI mouse behaviour detection system was implemented as a reproducible two-service Docker deployment consisting of (i) a custom processing service for video ingestion, preprocessing, and export, and (ii) Label Studio as the annotation interface. The processor automatically registered videos, embedded experimental metadata, generated annotation tasks via the Label Studio API, and exported structured mouse-level behavioural datasets.

*Video preprocessing and performance optimization*

Because mice were recorded in fixed cage positions, regions of interest were defined once and applied across videos to generate per-mouse clips using FFmpeg. To improve annotation efficiency, lightweight proxy videos were generated for responsive playback. Optional motion-based filtering was implemented to flag active segments but was used only as a supplementary aid due to high baseline activity levels.

*Annotation workflows and dataset generation*

Both multi-mouse and single-mouse annotation configurations were supported to maximize annotation efficiency and dataset flexibility. Metadata, including mouse identity and treatment group, were embedded at import and preserved during export. This pipeline enabled efficient, blinded identification and quantification of nasal rubbing events and produced structured datasets suitable for downstream behavioural analysis and future machine learning applications.

*Code availability*

The video analysis and annotation pipeline developed for this study is available as an open-source, containerized software package. The tool enables reproducible preprocessing, annotation, and export of structured behavioral datasets from multi-mouse video recordings. The complete source code, deployment configuration, annotation schemas, and documentation are available at: https://zenodo.org/records/19257529

#### Statistical Analysis

Statistical analyses were performed using Graph Pad Prism Version 10. Data are displayed as mean ± SEM. Statistical significance was determined using unpaired Student's t test, one or two-way ANOVA, as appropriate, and indicated in the figure legends. Significance is represented by the following scheme: *$p \leq 0.05$, **$p<0.01$, ***$p<0.001$, ****$p<0.0001$.

**Table S1. Overview of murine models of cat-AR.**

| reference | sensitization | | | challenge | | |
|---|---|---|---|---|---|---|
| | *route* | *dose (volume)* | *schedule* | *route* | *dose (volume)* | *schedule* |
| Zhu et al. 2005[S](19) | IP<br>IT (boost) | 5 µg Fel d 1 (NS)<br>1 µg Fel d 1 (NS) | 2x, 13 days apart<br>4x, 1-2 days apart | IT | 1 ug Fel d 1 (NS) | 1x |
| Terada et al. 2006[S](20) | IP<br>IT (boost) | 5 µg purified natural Fel d 1 in alum (160 µL)<br>1 µg native Fel d 1 (NS) | 2x, 14 days apart<br>4x, 1-3 days apart | IT | 1 ug native Fel d 1 (NS) | 3x, 7 days apart |
| | IP<br>IT (boost) | 1 or 10 µg purified natural Fel d 1 in alum (160 µL)<br>1 µg Fel d 1 (NS) | 1x<br>1x | IT | 1 µg Fel d 1 (NS) | 2x, 14 days apart |
| Neimert-Andersson et al. 2008[S](21) | SC | 1 µg recombinant Fel d 1 with alum (NS) | 3x, 14 days apart | IN | 10 µg cat dander extract (20 µL) | daily for 3 days |
| Thunberg et al. 2009[S](22) | SC | 1 µg recombinant Fel d 1 with alum (NS) | 3x, 14 days apart | IN | 10 µg cat dander extract (~20 µL) | daily for 3 days |
| Schmitz et al. (2009) (23) | IP | 1 µg natural Fel d 1 with alum (~200 µL) or 1 µg Fel d 1 and ovalbumin with alum (~200 µL) | 1x | IP | 10 µg recombinant Fel d 1 in PBS (~300 µL) | 1x |
| Saarne et al. 2011[S](24) | SC | 1 µg recombinant Fel d 1 with alum (NS) | 3x, 14 days apart | IN | 10 µg cat dander extract (20 µL) | daily for 3 days |
| Grundström et al. 2012[S](25) | SC | 1 µg recombinant Fel d 1 with alum in PBS (~200 µL) | 3x, 14 days apart | IN | 10 µg cat dander extract (20 µL) | daily for 3 days |
| Grundström et al. 2015[S](26) | SC | 1 µg recombinant Fel d 1 with alum in PBS (~200 µL) | once a week for 2 weeks | IN | 50 µg cat dander extract with 1 µg recombinant Fel d 1 (25 µL) | 3 consecutive days/week for a total of 7 weeks |
| Tasaniyananda et al. 2016[S](27) | IP | crude cat extract in PBS containing 10 µg natural Fel d 1 with alum (200 µL) | once a week for 3 weeks | IN (days 21-27)<br>Nebulized (days 34-36) | crude cat extract in PBS containing 1 µg Fel d 1 (20 µL; 10 µL per nostril)<br>10 mg crude cat hair extract in PBS (NS) | daily for 7 days<br>daily for 3 days |
| Mai et al. 2025[S](28) | IP | 1 mg/mL recombinant Fel d 1 with alum (100 µL) | once a week for 2 weeks | IT | 1 mg/mL recombinant Fel d 1 (50 µL) | 3x, 2 days apart |

IP = intraperitoneal; IN = intranasal; SC = subcutaneous; IT = intratracheal; NS = not specified

**Table S2. Fel d 1 concentrations in air samples from experimental and natural exposures.**

| reference | sampling location | concentration |
|---|---|---|
| **experimental exposures** | | |
| Luczynska et al. (1990) (29) | cat vivarium | 40 ng/$m^3$ |
| Wood et al. (1993) (30) | cat challenge room with live cats | 13 to 37967 ng/$m^3$ |
| Bollinger et al. (1996) (31) | cat challenge model (low-level) | 13.1 to 448 ng/$m^3$ |
| | cat challenge model (high-level) | 622 to 31381 ng/$^3$ |
| Larson et al. (2020) (32) | environmental exposure chamber | 10 to 500 ng/$m^3$ |
| Yang et al. (2022) (33) | naturalistic exposure chamber | 15.4 to 167.5 ng/$m^3$ |
| Haya et al. (2025) (34) | EnviroMini | 23 to 85 ng/$m^3$ |
| | naturalistic exposure chamber | 24 to 124 ng/$m^3$ |
| **natural exposures** | | |
| Luczynska et al. (1990) (29) | houses with cats | 2-20 ng/$m^3$ |
| | houses without cats | <0.2 ng/$m^3$ |
| Wood et al. (1993) (30) | homes with cats | 2 to 468.5 ng/$m^3$ |
| Bollinger et al. (1996) (31) | homes with cats | 1.8 to 578 ng/$m^3$ |
| | homes without cats | 2.8 to 88.5 ng/m3 |
| Custovic et al. (1998) (35) | public spaces | 0.09 to 0.22 ng/$m^3$ |
| Custovic et al. (1999) (36) | homes with cats | 0.4 to 22.3 ng/$m^3$ |
| | homes without pets | 0.16 to 1.8 ng/$m^3$ |
| Almqvist et al. (1999) (37) | classrooms with many cat owners | 2.94 ng/$m^3$ |
| | classrooms with few cat owners | 0.59 ng/$m^3$ |
| | homes without cats | 0.15 ng/$m^3$ |
| Almqvist et al. (1999) (37) | homes with a cat | 19.8 ng/$m^3$ |
| Niesler et al. (2016) (38) | homes with indoor cat | 0.09 to 2.21 ng/$m^3$ |
| | homes with outdoor cat | 0.07 to 0.1 ng/$m^3$ |

**Table S3. Time chart of clinical study procedures.**

| timepoints (hours) | symptom collection | blood sampling | nasal sampling |
|---|---|---|---|
| 0 | ✓ | ✓ | ✓ |
| 0.25 | ✓ | | |
| 0.5 | ✓ | | |
| 1 | ✓ | | |
| 1.5 | ✓ | | |
| 2 | ✓ | | |
| 2.5 | ✓ | | |
| 3 | ✓ | ✓ | ✓ |
| 4 | ✓ | | |
| 5 | ✓ | | |
| 6 | ✓ | ✓ | |
| 7 | ✓ | | |
| 8 | ✓ | | |
| 9 | ✓ | | |
| 10 | ✓ | | |
| 11 | ✓ | | |
| 12 | ✓ | | |
| 24 | ✓ | ✓ | ✓ |

**Table S4. Scoring of nasal symptoms in clinical study.**

| symptom | 0 | 1 | 2 | 3 |
|---|---|---|---|---|
| sneezing (/3) | symptom is completely absent | symptom is present, but not bothersome | symptom is bothersome, but tolerable | symptom is hard to tolerate, desiring treatment |
| runny nose/ post-nasal drip (/3) | | | | |
| nasal congestion/ stuffiness (/3) | | | | |
| itchy nose (/3) | | | | |

**Table S5. Criteria for appearance and activity scoring for health monitoring of mice.**

| appearance criteria | score | | Activity Criteria | Score |
|---|---|---|---|---|
| normal (hair coat and posture, eyes open) | 0 | | bright, alert, and responsive | 0 |
| slightly hunched, eyes partially closed | 1 | | active but hunched posture | 1 |
| moderately hunched, eyes moderately closed, pale mucous membranes | 2 | | less active when observed outside of cage but active when stimulated | 2 |
| ruffled coat, very hunched, eyes completely closed, pale mucous membranes | 3 | | inactive even when simulated (no response to touch, no righting reflex) | 3 |

**Table S6. Comparison of human and mouse models.**

| comparator | mouse model | human model |
|---|---|---|
| **experimental setup** | | |
| allergen type | cat dander | cat dander |
| allergen preparation | solution | powder |
| route of exposure | intranasal | inhalation |
| average Fel d 1 administration | 54 ng | 54 ng |
| **immunological effects** | | |
| nasal symptoms | ↑↑↑ | ↑↑↑ |
| systemic testing: | | |
| location | ear | forearm |
| positivity | ✓ | ✓ |
| serum total IgE | | |
| 3 hours | ↑ | ↑ |
| 6 hours | ↓ | – |
| serum sIgE | | |
| 3 hours | n/a | ↑ |
| 6 hours | n/a | ↓ |
| nasal total IgE | | |
| 3 hours | n/a | ↑ |
| 24 hours | n/a | ↓ |
| nasal sIgE | | |
| 3 hours | n/a | ↑ |
| 6 hours | n/a | – |
| 24 hours | n/a | – |
| nasal eosinophils | | |
| 3 hours | ↑ | ↑↑ |
| 6 hours | ↑↑ | ↑ |
| CD11b$^{int}$ MHC II$^{int–hi}$ eosinophils | | |
| 3 hours | ↓ | n/a |
| 6 hours | ↑ | n/a |
| CD11b$^{lo}$ MHC II$^{int–hi}$ eosinophils | | |
| 3 hours | ↓ | n/a |
| 6 hours | ↑ | n/a |
| nasal localization | | |
| CD45$^{+}$ immune cells | ↑ Levels II and III | n/a |
| EPX | ↑ Level III | n/a |
| PAS | – | n/a |
| epithelial thickness | (↑) Level III | n/a |
| aerated space | (↓) Level III | n/a |

n/a = not applicable

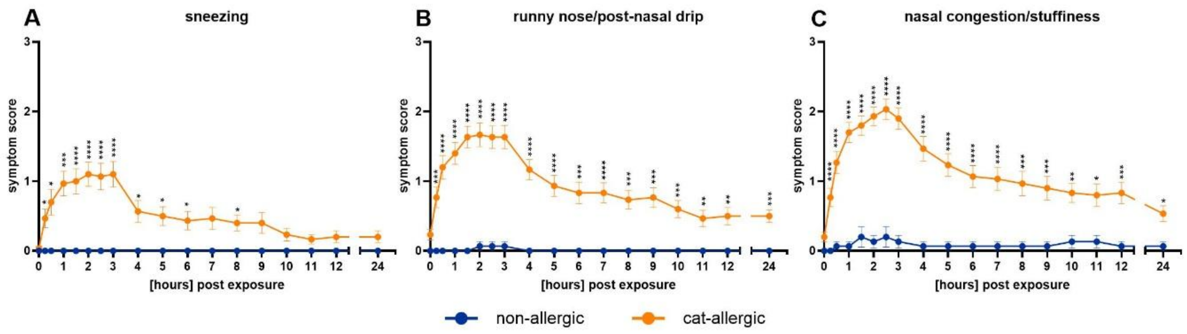


**Fig S1. Nasal symptoms after human experimental cat dander exposure.**

Cat-allergic and non-allergic participants were exposed to airborne cat dander in the SPaC-EEU for 3 hours. Symptom scores were captured on paper diary cards, with participants ranking them on a scale from 0 to 3. Symptom scores of **(A)** sneezing, **(B)** runny nose/post-nasal drip, and **(C)** nasal congestion/stuffiness. n = 15-31 participants/group. Mixed-effects analysis with Sidak's multiple comparisons test. *p≤0.05, **p<0.01, ***p<0.001, ****p<0.0001.

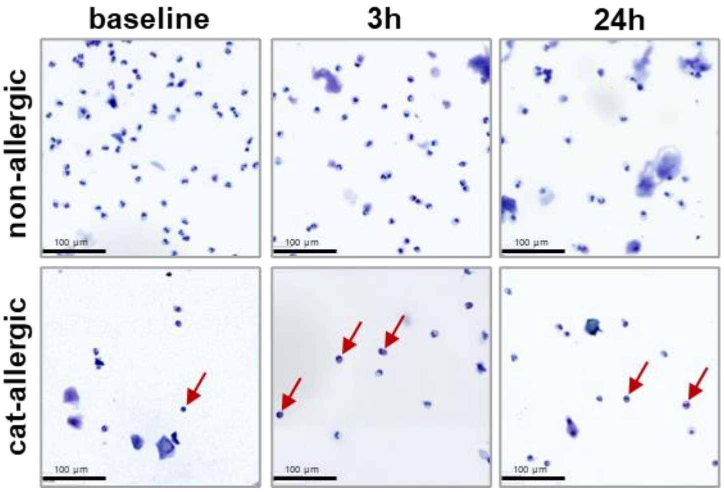


**Fig S2. Eosinophil detection in human nasal lavage samples.**

Representative images for Cytospin analysis of nasal lavage samples at baseline, 3 hours, and 24 hours post cat dander exposure in SPaC-EEU. Arrows highlight eosinophils.

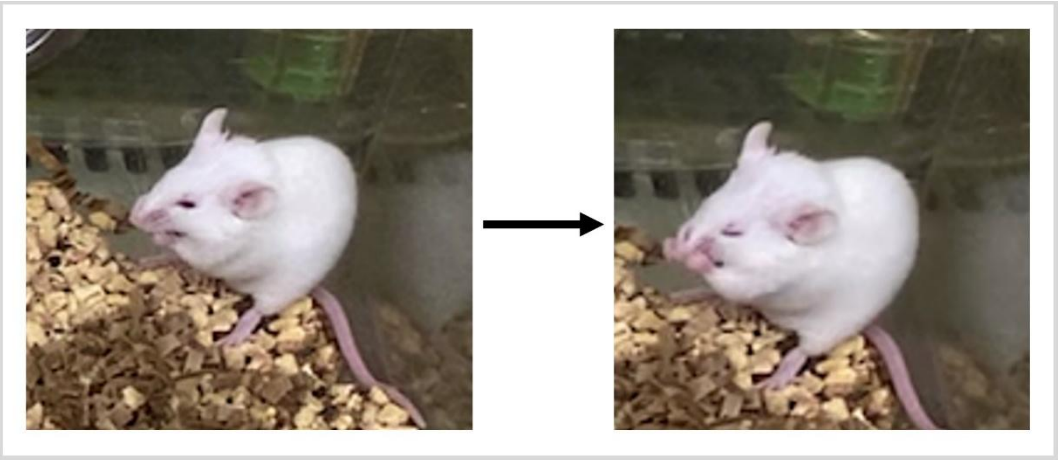

**Fig S3. Example of nasal rubbing in mice.**

Mice were video recorded for 30 minutes after each sensitization or challenge to detect nasal rubbing. Potential rubbing frames were segregated using a code, and occurrences of rubbing episodes and duration of nasal rubbing episodes were captured and tallied. The images show an example of nasal rubbing.

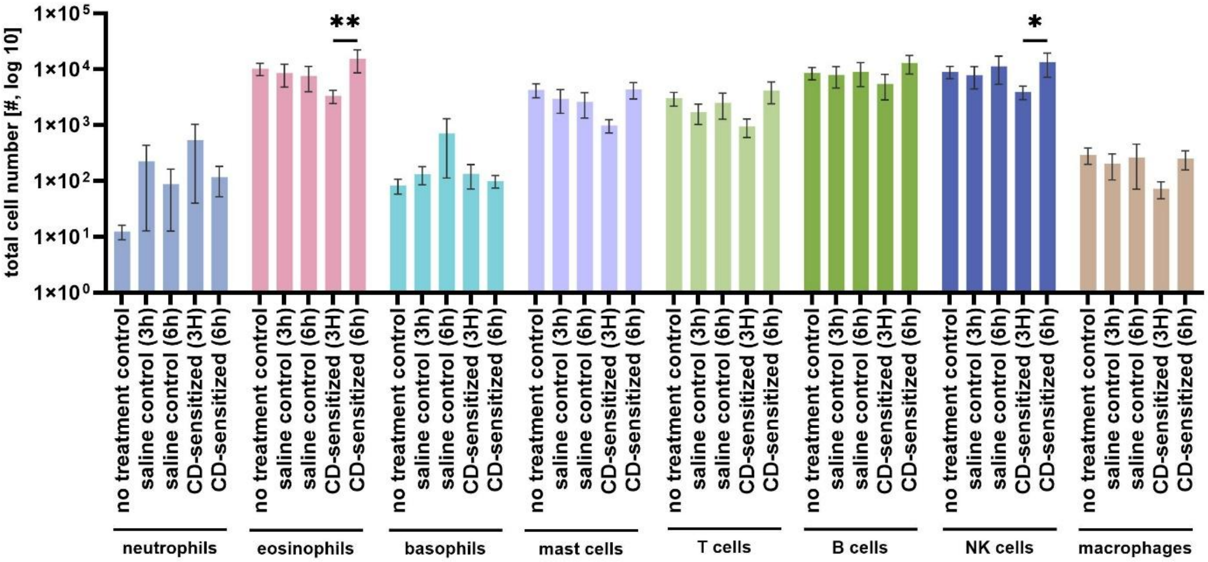


**Fig S4. Total numbers of nasal immune cells in CD-sensitized mice.**

Nasal immune cells are shown as total cell numbers (log 10) across experimental groups. n = 8 mice/group. Two-way ANOVA with Tukey's multiple comparisons test. *p≤0.05, **p<0.01.

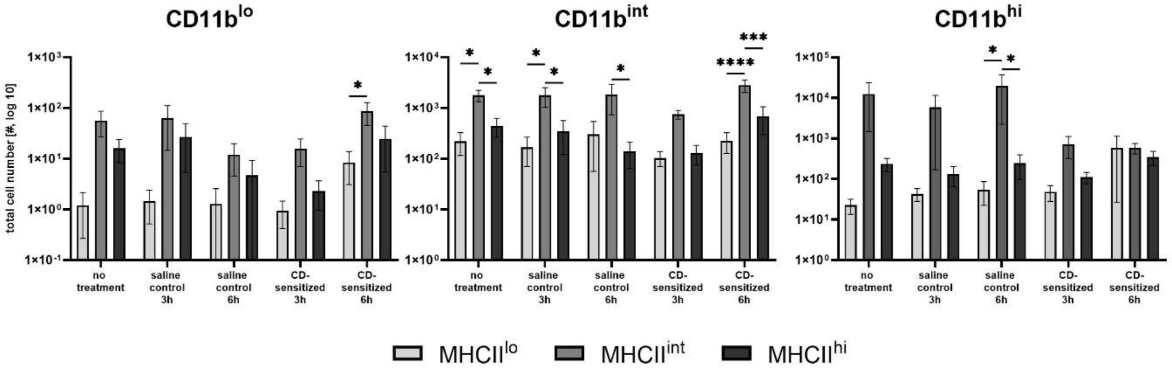


**Fig S5. Flowcytometric detection of CD11b and MHC II expression on murine nasal eosinophils.**

Eosinophils (CD45$^+$ Ly6G$^-$ F4/80$^{int_hi}$ IL5R$^+$) from nasal lavage were subcategorized in low or intermediate CD11b expression as well as low, intermediate, or high MHC II expression. n=8/group. Two-way ANOVA with Tukey's multiple comparisons test. *p≤0.05, ***p<0.001, ****p<0.0001.

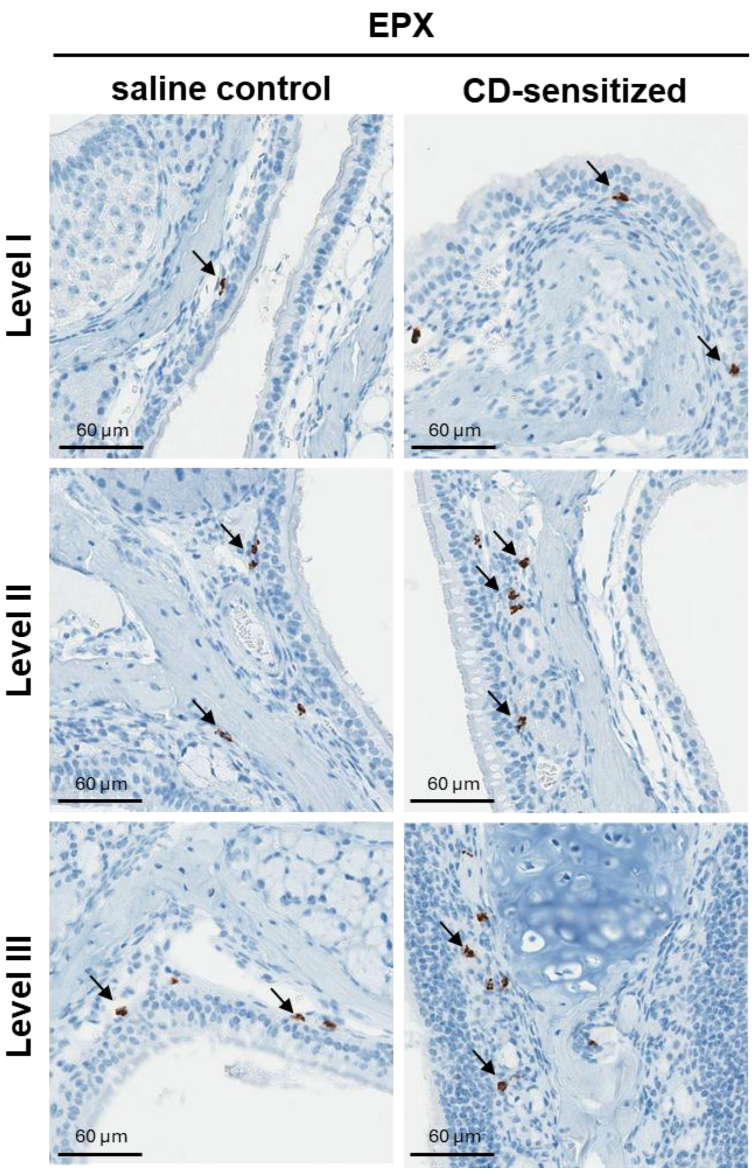


**Fig S6. Nasal localization of EPX in mice.**

Representative immunohistochemical staining images in 40x magnification. Arrows indicate representative EPX staining.

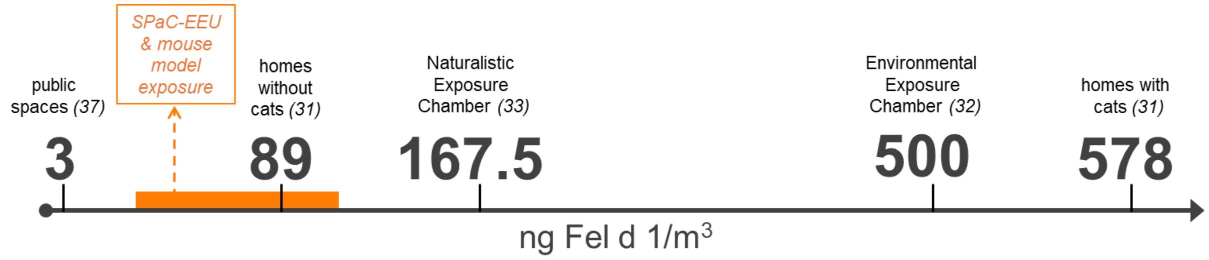


**Fig S7. Visual representation of Fel d 1 exposure concentrations.**

Clinical and mouse exposure models described in this manuscript employed Fel d 1 exposures at concentrations consistent with the lower to mid-range of environmental Fel d 1 allergen exposure.

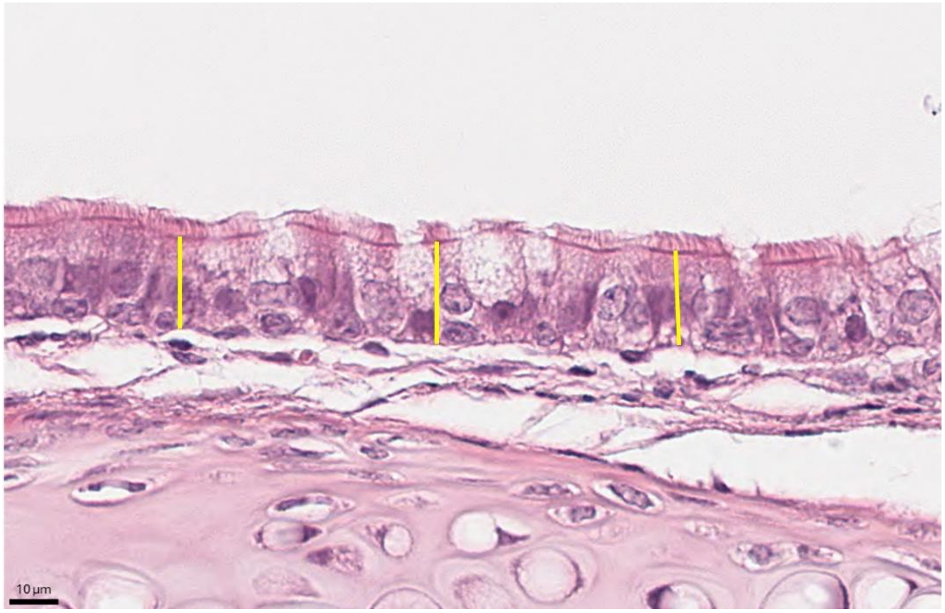


**Fig S8. Measurement of epithelial thickness in the murine nasal mucosa.**

Example for the measurement of epithelial thickness in a region of interest (ROI, perpendicular yellow lines) in the respiratory mucosa of mice. H&E staining, 400x magnification.

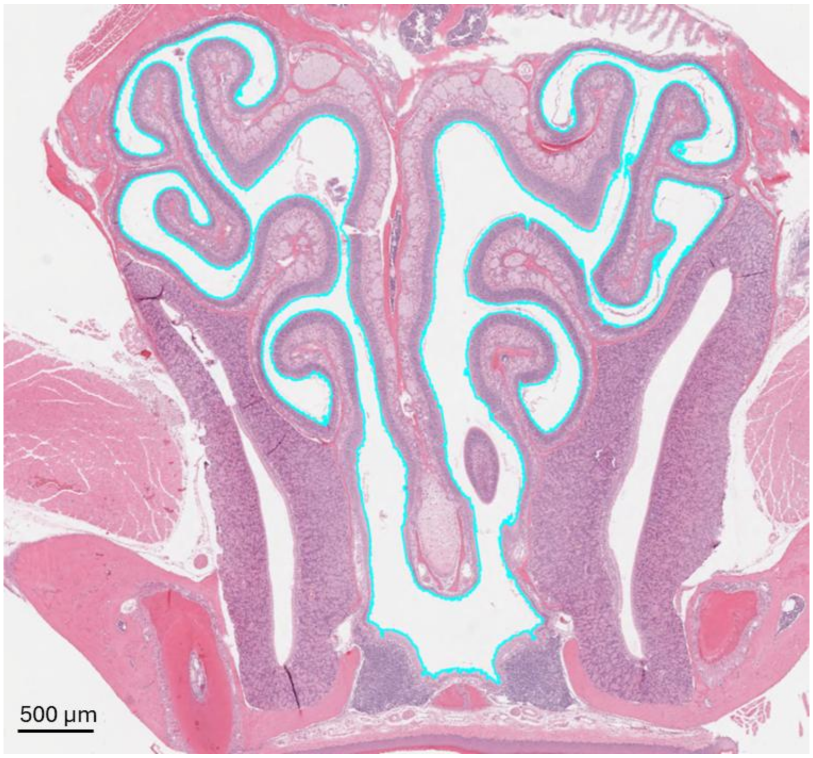


**Fig S9. Annotation of the aerated area at Level III of the mouse nose.**

The aerated area was quantified at nasal coronal level III of the mouse nose using the Create Thresholder function. This approach generated a contiguous annotation of the aerated compartment (as shown with the blue outline above), which was quantified as total aerated area in $\mu m^2$. H&E staining, 10x magnification.

#### Supplemental references


1. Hossenbaccus L, Walker T, Ellis AK. Technical validation of controlled exposure to cat dander in the specialized particulate control environmental exposure unit (SPaC-EEU). Allergy Asthma Clin Immunol. 2025;21(1):6.
2. Hossenbaccus L, Linton S, Garvey S, Botting H, Walker T, Steacy L, et al. Clinical validation of controlled exposure to cat dander in the Specialized Particulate Control Environmental Exposure Unit (SPaC-EEU). Allergy Asthma Clin Immunol. 2025;21(1):33.
3. Hossenbaccus L, Linton S, Thiele J, Steacy L, Walker T, Malone C, et al. Clinical validation of controlled exposure to house dust mite in the environmental exposure unit (EEU). Allergy Asthma Clin Immunol. 2021;17(1):34.
4. Pfaar O, Calderon MA, Andrews CP, Angjeli E, Bergmann KC, Bonlokke JH, et al. Allergen exposure chambers: harmonizing current concepts and projecting the needs for the future - an EAACI Position Paper. Allergy. 2017;72(7):1035–42.
5. Pfaar O, Bergmann KC, Bonini S, Compalati E, Domis N, de Blay F, et al. Technical standards in allergen exposure chambers worldwide - an EAACI Task Force Report. Allergy. 2021;76(12):3589–612.
6. Hossenbaccus L, Linton S, Thiele J, Steacy L, Walker T, Malone C, et al. Biologic Responses to House Dust Mite Exposure in the Environmental Exposure Unit. Front Allergy. 2021;2:807208.
7. Walker TJ, Steacy LM, Ellis AK. Preliminary Proof of House Dust Mite Distribution Capability in the Environmental Exposure Unit. Journal of Allergy and Clinical Immunology. 2017;139(2):AB119–AB.
8. Rawls M, Thiele J, Adams DE, Steacy LM, Ellis AK. Clinical symptoms and biomarkers of Bermuda grass-induced allergic rhinitis using the nasal allergen challenge model. Ann Allergy Asthma Immunol. 2020;124(6):608–15 e2.
9. Linton S, Hossenbaccus L, Davis A, Thiele J, Garvey S, Botting H, et al. Characterizing the symptomatology and pathophysiology of allergic rhinitis using a nasal allergen challenge model - a subset of the allergic rhinitis microbiome study. Allergy Asthma Clin Immunol. 2025;21(1):36.
10. Becker KJ. Strain-Related Differences in the Immune Response: Relevance to Human Stroke. Transl Stroke Res. 2016;7(4):303–12.
11. ICH Harmonized Guideline: Detection of Reproductive and Developmental Toxicity for Human Pharmaceuticals. https://database.ich.org/sites/default/files/S5-R3_Step4_Guideline_2020_0218.pdf. Accessed on February 16, 2026.
12. Liu W, Yu Z, Wang Z, Waubant EL, Zhai S, Benet LZ. Using an animal model to predict the effective human dose for oral multiple sclerosis drugs. Clin Transl Sci. 2023;16(3):467–77.
13. Phillips JE. Inhaled efficacious dose translation from rodent to human: A retrospective analysis of clinical standards for respiratory diseases. Pharmacol Ther. 2017;178:141–7.
14. Rodrigues A, Gualdi LP, de Souza RG, Vargas MH, Nunez NK, da Cunha AA, et al. Bacterial extract (OM-85) with human-equivalent doses does not inhibit the development of asthma in a murine model. Allergol Immunopathol (Madr). 2016;44(6):504–11.
15. Leekha A, Saeedi A, Kumar M, Sefat K, Martinez-Paniagua M, Meng H, et al. An intranasal nanoparticle STING agonist protects against respiratory viruses in animal models. Nat Commun. 2024;15(1):6053.
16. National Institute of Environmental Health Sciences. Nasal Cavity: Revised Guides for Organ Sampling and Trimming in Rats and Mice.
17. Bankhead P, Loughrey MB, Fernandez JA, Dombrowski Y, McArt DG, Dunne PD, et al. QuPath: Open source software for digital pathology image analysis. Sci Rep. 2017;7(1):16878.
18. Sanchez-Montalvo A, Lecocq M, Bouillet E, Steelant B, Gohy S, Froidure A, et al. Validation and shortcomings of the most common mouse model of chronic rhinosinusitis with nasal polyps. Rhinology. 2024;62(4):446–56.
19. Zhu D, Kepley CL, Zhang K, Terada T, Yamada T, Saxon A. A chimeric human-cat fusion protein blocks cat-induced allergy. Nat Med. 2005;11(4):446–9.
20. Terada T, Zhang K, Belperio J, Londhe V, Saxon A. A chimeric human-cat Fcgamma-Fel d1 fusion protein inhibits systemic, pulmonary, and cutaneous allergic reactivity to intratracheal challenge in mice sensitized to Fel d1, the major cat allergen. Clin Immunol. 2006;120(1):45–56.
21. Neimert-Andersson T, Thunberg S, Swedin L, Wiedermann U, Jacobsson-Ekman G, Dahlen SE, et al. Carbohydrate-based particles reduce allergic inflammation in a mouse model for cat allergy. Allergy. 2008;63(5):518–26.

22. Thunberg S, Neimert-Andersson T, Cheng Q, Wermeling F, Bergstrom U, Swedin L, et al. Prolonged antigen-exposure with carbohydrate particle based vaccination prevents allergic immune responses in sensitized mice. Allergy. 2009;64(6):919–26.
23. Schmitz N, Dietmeier K, Bauer M, Maudrich M, Utzinger S, Muntwiler S, et al. Displaying Fel d1 on virus-like particles prevents reactogenicity despite greatly enhanced immunogenicity: a novel therapy for cat allergy. J Exp Med. 2009;206(9):1941–55.
24. Saarne T, Neimert-Andersson T, Gronlund H, Jutel M, Gafvelin G, van Hage M. Treatment with a Fel d 1 hypoallergen reduces allergic responses in a mouse model for cat allergy. Allergy. 2011;66(2):255–63.
25. Grundstrom J, Neimert-Andersson T, Kemi C, Nilsson OB, Saarne T, Andersson M, et al. Covalent coupling of vitamin D3 to the major cat allergen Fel d 1 improves the effects of allergen-specific immunotherapy in a mouse model for cat allergy. Int Arch Allergy Immunol. 2012;157(2):136–46.
26. Grundstrom J, Saarne T, Kemi C, Gregory JA, Waden K, Pils MC, et al. Development of a mouse model for chronic cat allergen-induced asthma. Int Arch Allergy Immunol. 2014;165(3):195–205.
27. Tasaniyananda N, Chaisri U, Tungtrongchitr A, Chaicumpa W, Sookrung N. Mouse Model of Cat Allergic Rhinitis and Intranasal Liposome-Adjuvanted Refined Fel d 1 Vaccine. PLoS One. 2016;11(3):e0150463.
28. Mai Y, Sun X, Liu X, Chen H, Liang X, Huang Y, et al. Studies on nanoprotein vaccine alleviating symptoms of mice allergic to rFel d 1. Front Immunol. 2025;16:1524929.
29. Luczynska CM, Li Y, Chapman MD, Platts-Mills TA. Airborne concentrations and particle size distribution of allergen derived from domestic cats (Felis domesticus). Measurements using cascade impactor, liquid impinger, and a two-site monoclonal antibody assay for Fel d I. Am Rev Respir Dis. 1990;141(2):361–7.
30. Wood RA, Laheri AN, Eggleston PA. The aerodynamic characteristics of cat allergen. Clin Exp Allergy. 1993;23(9):733–9.
31. Bollinger ME, Eggleston PA, Flanagan E, Wood RA. Cat antigen in homes with and without cats may induce allergic symptoms. J Allergy Clin Immunol. 1996;97(4):907–14.
32. Larson D, Patel P, Salapatek A, Couroux P, Whitehouse D, Pina A, et al. Nasal allergen challenge and environmental exposure chamber challenge: A randomized trial comparing clinical and biological responses to cat allergen. The Journal of allergy and clinical immunology. 2020;145(6).
33. Yang WH, Kelly S, Haya L, Mehri R, Ramesh D, DeVeaux M, et al. Cat allergen exposure in a naturalistic exposure chamber: A prospective observational study in cat-allergic subjects. Clin Exp Allergy. 2022;52(2):265–75.
34. Haya L, Kelly S, Van de Mosselaer S, Friedrich R, Yang J, Yang WH. Clinical Response to Cat Allergen in a Mobile Compared to a Fixed Naturalistic Exposure Chamber. Int Arch Allergy Immunol. 2025;186(12):1119–28.
35. Custovic A, Fletcher A, Pickering CA, Francis HC, Green R, Smith A, et al. Domestic allergens in public places III: house dust mite, cat, dog and cockroach allergens in British hospitals. Clin Exp Allergy. 1998;28(1):53–9.
36. Custovic A, Simpson B, Simpson A, Hallam C, Craven M, Woodcock A. Relationship between mite, cat, and dog allergens in reservoir dust and ambient air. Allergy. 1999;54(6):612–6.
37. Almqvist C, Larsson PH, Egmar AC, Hedren M, Malmberg P, Wickman M. School as a risk environment for children allergic to cats and a site for transfer of cat allergen to homes. J Allergy Clin Immunol. 1999;103(6):1012–7.
38. Niesler A, Scigala G, Ludzen-Izbinska B. Cat (Fel d 1) and dog (Can f 1) allergen levels in cars, dwellings and schools. Aerobiologia (Bologna). 2016;32(3):571–80.

#### GitHub Repository README

##### Mouse behavior annotation tool

Local-first, containerised video annotation platform for quantifying clinically relevant behavioural patterns in mice, with structured dataset export for downstream analysis and machine learning.

##### Overview

This tool was developed to support behavioural quantification of nasal rubbing events in mice exposed to inhaled allergens. The platform accelerates manual behavioural annotation while producing structured, reusable datasets suitable for downstream statistical analysis and future machine learning model development.

The system is designed for reproducibility, accessibility, and extensibility. It runs entirely locally using Docker containers, requires no Python installation on the host system, and provides a browser-based interface for efficient annotation.

Primary objectives:

1. Accelerate manual annotation of clinically relevant behaviour
2. Generate structured, mouse-level datasets suitable for future automated behavioural classification

##### Scientific context

Manual behavioural annotation of laboratory mice from continuous video recordings is time-consuming and prone to inefficiencies due to:

- large video file sizes
- multi-animal recordings requiring sequential review per mouse
- web playback latency
- lack of structured export formats linking annotations to experimental metadata

This tool addresses these challenges by:

- isolating mouse-specific video regions via fixed ROI cropping
- generating web-optimised proxy media
- embedding experimental metadata (mouse IDs, treatment groups) at task import
- exporting analysis-ready CSV datasets with mouse-resolved behavioural events

Importantly, this tool does NOT perform automated behavioural classification. It supports efficient manual annotation and dataset generation for downstream modelling.

##### System architecture

The platform uses a two-service architecture orchestrated via Docker Compose.

**Components:**

**1. Processor service**

Custom web dashboard responsible for:

- Video ingestion and metadata registration
- metadata parsing and registration via filename conventions and `mouse_map.csv` lookup
- ROI definition and cropping
- proxy media generation
- Label Studio project creation and task import via REST API
- annotation export and CSV dataset formatting

**Technology stack:**

Python: 3.9
Framework: Streamlit 1.29.0
FFmpeg: 7.1.3 System package (Debian)
Docker base image: python:3.9-slim

**2. Annotation service**

Label Studio provides annotation interface.
Label Studio version: 1.12.1
The processor communicates with Label Studio via REST API.

**Key API functions:**

- Create/update project with XML label configuration
- Import video tasks with embedded metadata
- Create local file storage connections
- Retrieve annotation results via snapshot export

**Authentication:**

Auto-retrieved API token from Label Studio SQLite database (see label_studio.py), with fallback to username/password session login via CSRF

**Data storage model**

A shared workspace directory serves as the primary data layer. The filesystem acts as the database, no SQL/NoSQL setup is required.

**Structure:**

```
workspace/
  raw/                              # Input videos (the "Inbox")
    {video_name}/                   # Per-video artifact folder (created at registration)
      {video_name}.json             # Metadata sidecar
      {video_name}_proxy.mp4        # Web-optimised video proxy
      {video_name}_audio.mp3        # Audio proxy for waveform rendering
```

```
  processed/            # Per-mouse cropped video clips + JSON sidecars
  outputs/              # Exported CSV datasets
  mouse_map.csv         # Group/treatment/mouse-ID lookup table
```

This structure ensures reproducibility and transparent dataset generation.

**Metadata system**

Each registered video has a JSON sidecar created during ingestion:

**Example:**

```
{
  "group": "Group 5",
  "treatment": "Control",
  "date": "Dec2",
  "mouse_ids": ["213696", "212754", "212274", "213665"],
  "original_file": "Control_Dec2-024.M4V",
  "full_path": "/workspace/raw/Control_Dec2-024.M4V"
}
```

Metadata is embedded into Label Studio tasks at import and preserved through the export pipeline.

**Metadata linkage hierarchy:**

1. Filename parsing
2. mouse_map.csv lookup
3. Folder structure fallback

**Region-of-Interest cropping**

Because animals occupy fixed cage positions, object detection is unnecessary. Instead, a "draw once, crop all" approach is used.

Workflow:

1. User defines 4 ROIs on first frame via an interactive canvas (streamlit-drawable-canvas)
2. Processor applies ROI coordinates to entire video
3. Per-mouse video clips generated using FFmpeg with parallel processing (ThreadPoolExecutor)

**Benefits:**

• deterministic cropping
• reduced computational complexity
• no object tracking errors
• reusable training clips for future ML work

**FFmpeg cropping command:**

```
ffmpeg -i input.mp4 \
  -filter_complex "crop=w:h:x:y" \
  -c:v libx264 -c:a aac -preset veryfast -crf 23 \
output.mp4
```

Four crops are processed in parallel using `concurrent.futures.ThreadPoolExecutor`

**Proxy media generation**
Large video files slow browser playback and annotation.
To improve performance, lightweight proxy media are generated.
Video proxy:
Resolution: 720p
Codec: H.264
Bitrate: 26
Command: ffmpeg -y -i input -vf scale=-2:720 -c:v libx264 -crf 26 -preset veryfast -an -movflags +faststart output.mp4
Audio proxy:
Codec: AAC
Sample rate: 44100 Hz
Purpose:
• waveform rendering
• synchronized playback
Proxy generation improves responsiveness and annotation throughput.

**Annotation modes**
Two annotation modes are supported via separate Label Studio projects.

**Multi-mouse mode (primary workflow)**
Annotates the full cage view. The researcher labels all 4 mice in a single pass using positional labels: `Rubbing (M1)`, `Rubbing (M2)`, `Rubbing (M3)`, `Rubbing (M4)`.

**Advantages:**
• faster annotation
• preserves spatial context

**Single-mouse mode**
Annotates cropped mouse-specific clips.

**Advantages:**

• simplified annotation
• ideal for dataset construction

Both modes export mouse-resolved behavioural events.

**Export pipeline**

Annotations are exported as structured tabular data.

**Export formats:**

• CSV
• JSON

Export columns:

MouseID, Label_Name, Group, Date, Treatment, Behavior, Rub_Start_Time, Rub_End_Time, Duration_Seconds, Video_File, Annotator_ID, Task_ID, Status

**Example export:**

```
MouseID,Label_Name,Group,Date,Treatment,Behavior,Rub_Start_Time,Rub_End_Time,Duration_Seconds,Video_File,Annotator_ID,Task_ID,Status
213696,Mouse 1,Group 5,Dec2,Control,Rubbing,42.21,196.98,154.77,Control_Dec2-024_proxy.mp4,1,7,Submitted
212754,Mouse 2,Group 5,Dec2,Control,Rubbing,401.99,600.98,198.99,Control_Dec2-024_proxy.mp4,1,7,Submitted
```

**Dataset generation**

The system produces reusable structured datasets.

Output includes:

• per-mouse behavioural annotations
• aligned metadata
• reusable cropped video clips

This enables downstream machine learning development.

Future extensions:

• automated classification
• behaviour prediction
• dataset expansion

**Installation**

Prerequisites:

Docker Desktop

Clone repository:

git clone https://github.com/Jackmactaylor/mouse-behaviour-analysis-tools

cd mouse-behavior-annotation

**Start system:**
docker compose up
Access interface:
Processor dashboard:
http://localhost:8501
Label Studio:
http://localhost:8080

Default Label Studio credentials: `` / `password123` (configurable in `docker-compose.yml`).

**Reproducibility**
The system ensures reproducibility via:
• containerised deployment
• version-controlled configuration
• structured data storage
• deterministic preprocessing pipeline

**Repository contents**

```
src/                     # Python application code
  app.py                 #   Streamlit dashboard (5 pages)
  setup_workspace.py     #   Workspace directory initializer
  components/      #   UI components (ROI selector, Label Studio XML configs)
  utils/                 #     Core logic (video processing, metadata, Label Studio API, export)
docker/                  # Dockerfile and container configuration
  processor/
    Dockerfile
    requirements.txt
docs/                    # Documentation, plans, and guides
workspace/               # Data layer (raw videos, processed clips, exports)
tests/                   # Validation and debugging scripts
docker-compose.yml       # Service orchestration
README.md                # This file
```

**Version information**
Python: 3.9
Docker: python:3.9-slim

Label Studio: 1.12.1
Streamlit: 1.29.0
FFmpeg: 7.1.3

**Intended use**

This tool is designed for:

• behavioural neuroscience
• immunology studies
• allergy and infection models
• translational animal research

**Citation**

If used in research, please cite:
[FILL IN MANUSCRIPT REFERENCE]

**Contact**

Joaquin Sanz
University of Zaragoza


**License**

MIT License. See LICENSE (https://github.com/JSanzR/mouse-behaviour-analysis-tools?tab=License-1-ov-file ) for details.
